# The neural mechanisms supporting the rise and fall of maternal aggression

**DOI:** 10.1101/2025.07.15.665001

**Authors:** Takashi Yamaguchi, Rongzhen Yan, Mashrur Khan, Kanishk Tewatia, Takuya Osakada, Srinivas Parthasarathy, Nirao M. Shah, Dayu Lin

**Author notes:** These authors contributed equally to this work (Takashi Yamaguchi, Rongzhen Yan).

## Abstract

To protect the helpless young, lactating females dramatically increase aggression towards intruders, known as maternal aggression. However, attack is costly and risky. When pups no longer exist, maternal aggression rapidly declines. Our study reveals the critical role of the pathway from posterior amygdala cells expressing estrogen receptor alpha (PA^Esr1^) to the ventrolateral part of ventromedial hypothalamus cells expressing neuropeptide Y receptor Y2 (VMHvl^Npy2r^) in the rise and fall of maternal aggression. Functional manipulations and recordings demonstrate that VMHvl-projecting PA^Esr1^ (PA^Esr1→VMHvl^) cells are naturally active and required for maternal aggression. During lactation, PA-VMHvl^Npy2r^ connection strengthens, and VMHvl^Npy2r^ excitability increases to facilitate attack. Furthermore, PA^Esr1^ expresses abundant oxytocin receptors, enabling oxytocin to boost PA^Esr1^ cell output. After pup separation, the oxytocin level drops, causing decreased maternal aggression, which can be restored by optogenetically increasing the oxytocin level. Thus, diverse forms of plasticity occur at the PA^Esr1^-VMHvl^Npy2r^ circuit to support need-based maternal aggression.

## Introduction

After birth, the young are vulnerable and helpless. To ensure their survival, mothers in nearly all mammalian species express a series of stereotypical behaviors to care for and protect the offspring. In mice, lactating females devote most of the day to nursing, crouching over and grooming pups, and retrieving the pups back to safety when they stray away accidentally. Additionally, when the pups are under the threat of a foreign intruder, mothers intensively attack the intruder to protect the young, a behavior known as maternal aggression^1–5^.

The estrogen receptor alpha-expressing cells in the ventrolateral part of the ventromedial hypothalamus (VMHvl^Esr1^) are essential for both male and female aggression^6–10^. In females, VMHvl^Esr1^ cells can be further separated into two topographically arranged subpopulations: the medially located VMHvl^Esr1^ cells (VMHvlm) are preferentially activated during maternal aggression, whereas the lateral ones (VMHvll) are mostly activated during female sexual behaviors^9,11^. A recent study identified Npy2r (VMHvl^Npy2r^) as an important molecular marker for the VMHvl aggression population in females^10^, whereas Npy2r negative VMHvl^Esr1^ cells (VMHvl^Npy2r-Esr1+^) are found relevant for mating^10,12,13^. We further showed that VMHvl^Npy2r-Esr1+^ cells overlap with VMHvl cholecystokinin A receptor (VMHvl^Cckar^) cells, a key population for female sexual receptivity^12,13^. During lactation, VMHvl^Npy2r^ cells, but not VMHvl^Npy2r-Esr1+^ or VMHvl^Cckar^ cells^10,13^, show higher *in vivo* responses to intruders compared to virgin females, although the cellular or synaptic changes underlying the response increase have not been investigated.

Beyond the VMHvl, the female aggression circuit remains largely unknown. Our previous study found that Esr1 cells in the posterior amygdala (PA^Esr1^), which are nearly all glutamatergic, provide strong excitatory inputs to the VMHvl to drive male aggression. Inhibiting PA^Esr1^ cells projecting to the VMHvl (PA^Esr1→VMHvl^) impairs aggression in male mice, whereas activating the cells has the opposite effect^14,15^. Furthermore, PA^Esr1^-VMHvl pathway is plastic. With repeated winning in male mice, PA^Esr1^-VMHvl connection potentiates, triggering additional changes in the local VMHvl network and VMHvl cell excitability, ultimately leading to heightened aggression^16,17^.

Given the critical role of the PA^Esr1^-VMHvl pathway in male aggression and experience-dependent behavior change, we investigated the potential involvement of this circuit in maternal aggression, especially whether plasticity in this circuit supports the dramatically increased aggression during motherhood. We found that PA^Esr1^-VMHvl circuit is also critical in maternal aggression as in male aggression. However, the neural mechanisms supporting the rise of aggression in lactating females are distinct from those in males. In particular, oxytocin-dependent modulation enables rapid changes in mothers’ aggression levels based on the need to protect the young^5,18^.

## Results

### Change in aggression level during the postpartum period

We previously found that maternal aggression varies greatly across strains of mice^9^. While 100% of lactating Swiss Webster (SW) females reliably attack juvenile intruders, only 60% of C57 females do so^1,9^. To increase the probability of observing maternal aggression, we crossed transgenic mice in C57 background with wildtype SW mice and used F1 generation as our experimental animals. Indeed, the hybrid females showed reliable maternal aggression. 83% of C57 × SW females attacked juvenile intruders on postpartum day (PPD) 1 (**Fig. 1a, b**). The aggression level increased on PPD3, peaked on PPD5, and stayed high until PPD7 (**Fig. 1c, d**). On PPD5, nearly all females attacked immediately and repeatedly upon intruder introduction (**Fig. 1c, d**). After the first postpartum week, aggression level gradually declined, and most females (5/6) stopped attacking the intruder by the end of the third week (**Fig. 1b-d**). Maternal aggression towards adult male and female intruders was much lower, as previously reported (**Extended Data Fig. 1a-d**)^19,20^. Thus, we used juveniles as intruders for all our subsequent experiments. The aggression of C57 × SW female mice is limited to the lactation period, as virgin females rarely attacked the intruders regardless of their estrous state (**Extended Data Fig. 1e**).

**Figure 1.**
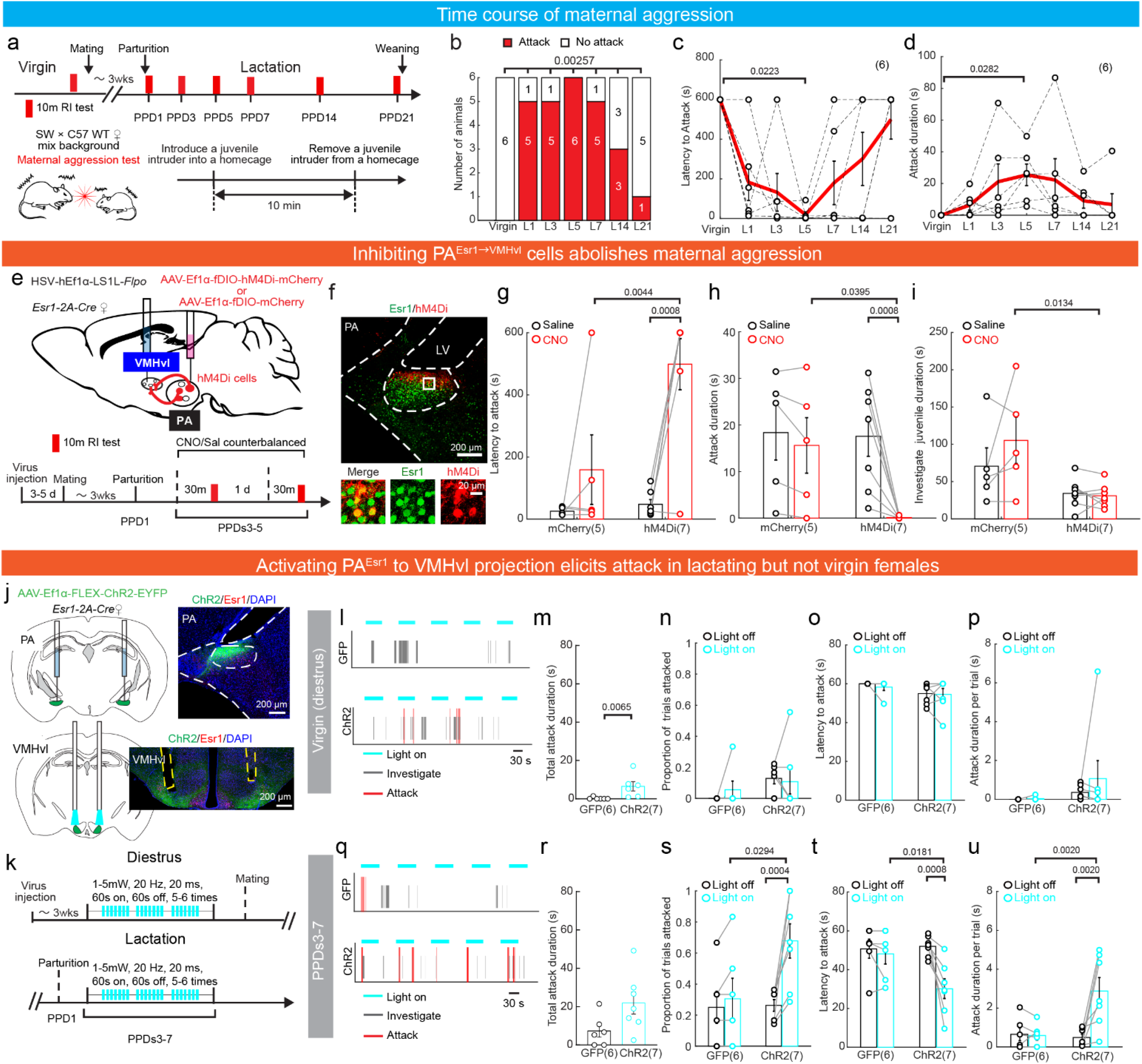
PA^Esr1^-VMHvl pathway bi-directionally modulates maternal aggression. **a**, Timeline to test aggression towards a juvenile intruder in C57×SW WT female mice across reproductive states. **b-d,** Fraction of females showing aggression **(b)**, their latency to attack **(c),** and total attack duration **(d)** at different reproductive states. **e,** Experimental schematics and test schedule. **f**, Histological images showing hM4Di expression and Esr1 staining in PA. The bottom shows enlarged views of the boxed areas; scale bars: 200 μm (top) and 20 μm (bottom). **g, h**, The latency to attack (**g**) increased, and durations of attack (**h**) significantly decreased after CNO injection in hM4Di but not mCherry group. For animals that showed no attack, the latency was 600 s. **i,** No change in juvenile investigation duration after CNO or saline injections. **j**, Experimental schematics and representative images showing ChR2-EYFP (green) and Esr1 (red) expression in PA and VMHvl. Blue, DAPI, red, Esr1; scale bar, 200 μm. **k**, Experimental timeline. **l, q**, Representative raster plots showing behaviors against juvenile intruders with and without light stimulation in virgin **(l)** and lactating **(q)** females. **m, r**, Total attack duration of ChR2 and GFP virgin diestrous **(m)** and lactating **(r)** females. **n-p**, No change in the proportion of trials the female attacked **(n)**, latency to attack from light onset **(o),** and attack duration per trial **(p)** during light-on and light-off periods in both ChR2 and GFP virgin diestrous females. **s-u**, In lactating ChR2 animals, but not GFP animals, the proportion of trials with attack increased **(s)**, the latency to attack decreased **(t),** and the average attack duration per trial increased **(u).** All bars or lines with error bars represent mean ± s.e.m. Numbers in parentheses represent the number of subject animals. Statistical significance was determined using **(b)** Cochran’s Q test followed by Mcnemar’s test with FDR correction of Benjamini and Hochberg, **(c, d)** Friedman test followed by Dunn’s multiple-comparison test, **(g-i, n-p, s-u)** two-way ANOVA with repeated measures followed by Bonferroni’s multiple-comparison test, **(m)** Mann Whitney test, and **(r)** unpaired t-test. All p values < 0.05 are indicated. See Supplementary Table 1 for more detailed statistics.

### PA^Esr1^**^→^**^VMHvl^ cells are necessary for maternal aggression

To investigate whether PA^Esr1→VMHvl^ cells are necessary for maternal aggression, we virally expressed hM4Di in the PA^Esr1→MHvl^ cells by injecting a retrograde HSV expressing Cre-dependent Flpo into the VMHvl and AAV-Ef1α-fDIO-hM4Di-mCherry (control: mCherry) into the PA of virgin Esr1-2A-Cre mice with C57 and SW mixed background (**Fig. 1e**). Histological analysis confirmed that the vast majority of hM4Di-mCherry-expressing cells were Esr1 positive in the PA (96.81 ± 0.03% cells from 3 animals; **Fig. 1f**). 3-5 days after virus injection, each female was paired with a male until becoming visibly pregnant. We then tested the female aggression level between PPDs 3 and 5, 30 minutes after either saline or CNO i.p. injection on separate days in a counterbalanced order (**Fig. 1e**). After saline injection, all females attacked the juveniles quickly and repeatedly (**Fig. 1g, h**). In contrast, only 2/7 hM4Di females attacked briefly during the 10-min tests after CNO injection, while all mCherry females continued to attack reliably (**Fig. 1g, h**). The decreased attack was not due to a loss of social interest in general, as the investigation duration did not change after PA^Esr1→VMHvl^ cell inactivation (**Fig. 1i**). In a separate experiment, we inhibited PA^Esr1^ cells as a whole and similarly found significantly reduced aggression in lactating females (**Extended Data Fig. 2a-d**). In contrast to maternal aggression, maternal care did not change after PA^Esr1→VMHvl^ or PA^Esr1^ cell inhibition in lactating females (**Extended Data Figs. 2e** and **3a, b**).

VMHvl also plays a critical role in female sexual behaviors^12,13,21^. Thus, we additionally asked whether PA^Esr1→VMHvl^ cells are necessary for female sexual receptivity, measured as the probability of lordosis, a back-arched posture to facilitate intercourse, and rejection^13,21^. Surprisingly, we found no difference in lordosis rate between saline- or CNO-injected days (**Extended Data Fig. 3a, c, d**). The males achieved intromission with similar latencies and durations when paired with CNO- or saline-injected test female mice (**Extended Data Fig. 3e-i**). Similar results were observed after inhibiting PA^Esr1^ cells, suggesting a non-critical role of PA^Esr1^ cells in female sexual behaviors (**Extended Data Fig. 2f-l**). Thus, PA^Esr1→VMHvl^ cells are specifically required for maternal aggression but not other social behaviors in females.

### PA^Esr1^-VMHvl projection can drive aggression in lactating females

To test whether PA^Esr1^ to VMHvl projection is sufficient to drive aggressive behaviors, we bilaterally injected AAV-Ef1α-FLEX-ChR2-EYFP into the PA and implanted the optic fibers above the VMHvl in female Esr1-2A-Cre mice with mixed C57 and SW background (**Fig. 1j**). Three weeks after virus injection, we first tested the stimulation-induced behavior changes in diestrous virgin female mice by delivering light pulses (470 nm, 20 Hz, 20 ms, 1.0 - 5.0 mW) for 60 s every 2-3 min after introducing a juvenile intruder (**Fig. 1k**). None of the diestrous females showed time-locked attack during PA^Esr1^ to VMHvl terminal activation, although ChR2 animals attacked more than GFP animals overall during the test session (**Fig. 1l-p**). After the aggression test with the juvenile intruder, we introduced the female into the home cage of a sexually experienced male and found light stimulation did not increase female sexual receptivity (**Extended Data Fig. 4a, b**). Both GFP and ChR2 animals consistently rejected the males with or without light stimulation, and the males never achieved intromission (**Extended Data Fig. 4b-d**).

After completing the tests in virgin females, we paired the females with males and tested the effect of PA^Esr1^ to VMHvl terminal activation on aggression again during PPDs 3 to 7 when the females were naturally aggressive. Upon light stimulation, ChR2 animals, but not GFP animals, significantly increased the probability of attacking the juvenile intruder (**Fig. 1q-u**). The latency to attack was lower, and the attack duration was higher during the light-on trials than during light-off trials (**Fig. 1t, u**). These results suggest that PA^Esr1^ is an important input to the VMHvl to drive female aggression. Its efficacy to induce attack is higher during lactation than virgin states.

### VMHvl^Npy2r^ cells increase responses corresponding to PA inputs during lactation

Why could PA inputs to the VMHvl drive time-locked attack in mothers but not in virgin females? To address this question, we investigated the VMHvl cell responses to PA inputs using *in vitro* patch-clamp recordings (**Fig. 2a**). We focused on VMHvl^Npy2r^ cells because of their reported central role in maternal aggression^10^. Indeed, as previously shown^10^, we found direct optogenetic activation of VMHvl^Npy2r^ cells reliably induced attack in virgin female mice, suggesting that the spiking activity of VMHvl^Npy2r^ cells plays a deterministic role in female attack initiation (**Extended Data Fig. 5**). We then performed current-clamp recordings of VMHvl^Npy2r^ cells and compared their spiking probability during PA-VMHvl terminal stimulation in virgin vs. lactating females (PPDs 3 - 5) (**Fig. 2b**). While PA terminal activation (5 ms, 20 Hz, 473 nm light pulses) induced action potentials only in <15% of VMHvl^Npy2r^ cells in virgin diestrous or estrous females, nearly half of (11/23) VMHvl^Npy2r^ cells spiked upon PA terminal activation in lactating females (**Fig. 2b, c**). The increased spiking probability of VMHvl^Npy2r^ in lactating females could be caused by changes in PA^Esr1^ to VMHvl connection strength or VMHvl cell excitability. To examine the former, we performed voltage-clamp recordings of VMHvl^Npy2r^ cells in virgins and mothers (PPDs 3-5) while optogenetically activating PA terminals to evoke excitatory and inhibitory postsynaptic currents (oEPSCs and oIPSCs) (**Fig. 2d, e**). As the case in males^15^, the oEPSC latency was short (mean ± s.e.m.: 3.49 ± 0.38 ms), indicating monosynaptic connection (**Fig. 2f**), whereas the latency of oIPSC was long (mean ± s.e.m.: 12.47 ± 1.31 ms), suggesting polysynaptic connection (**Fig. 2f**). Between virgin and lactating females, PA terminal stimulation is more likely to evoke EPSC of VMHvl^Npy2r^ cells in lactating females (79%, 19/24) than virgin females, although the percentage of cells responded to PA input in virgin females varied dramatically over the estrous cycle. During diestrus, 65% (13/20) of VMHvl^Npy2r^ cells showed optically evoked EPSC, while only 12% (2/16) did so during estrus (**Fig. 2e**). The oEPSC amplitude in estrous females also appeared lower than in diestrous and lactating females (**Fig. 2g**). This difference is likely not due to presynaptic changes as the pair-pulse ratio of oEPSCs is comparable across reproductive states (**Fig. 2h, i**). The probability of PA stimulation to evoke IPSC also varied with the reproductive state in a similar fashion as the oEPSC: high during lactation and low during estrus (**Fig. 2e, g**). This is not surprising as the oIPSC is likely secondary to excitatory transmission. Additionally, the spontaneous EPSC (sEPSC), but not IPSC (sIPSC), frequency of VMHvl^Npy2r^ cells was the lowest during estrus (**Fig. 2j-n**). These results suggest that the PA-VMHvl^Npy2r^ connection is the strongest during lactation and weakest during estrus.

**Figure 2.**
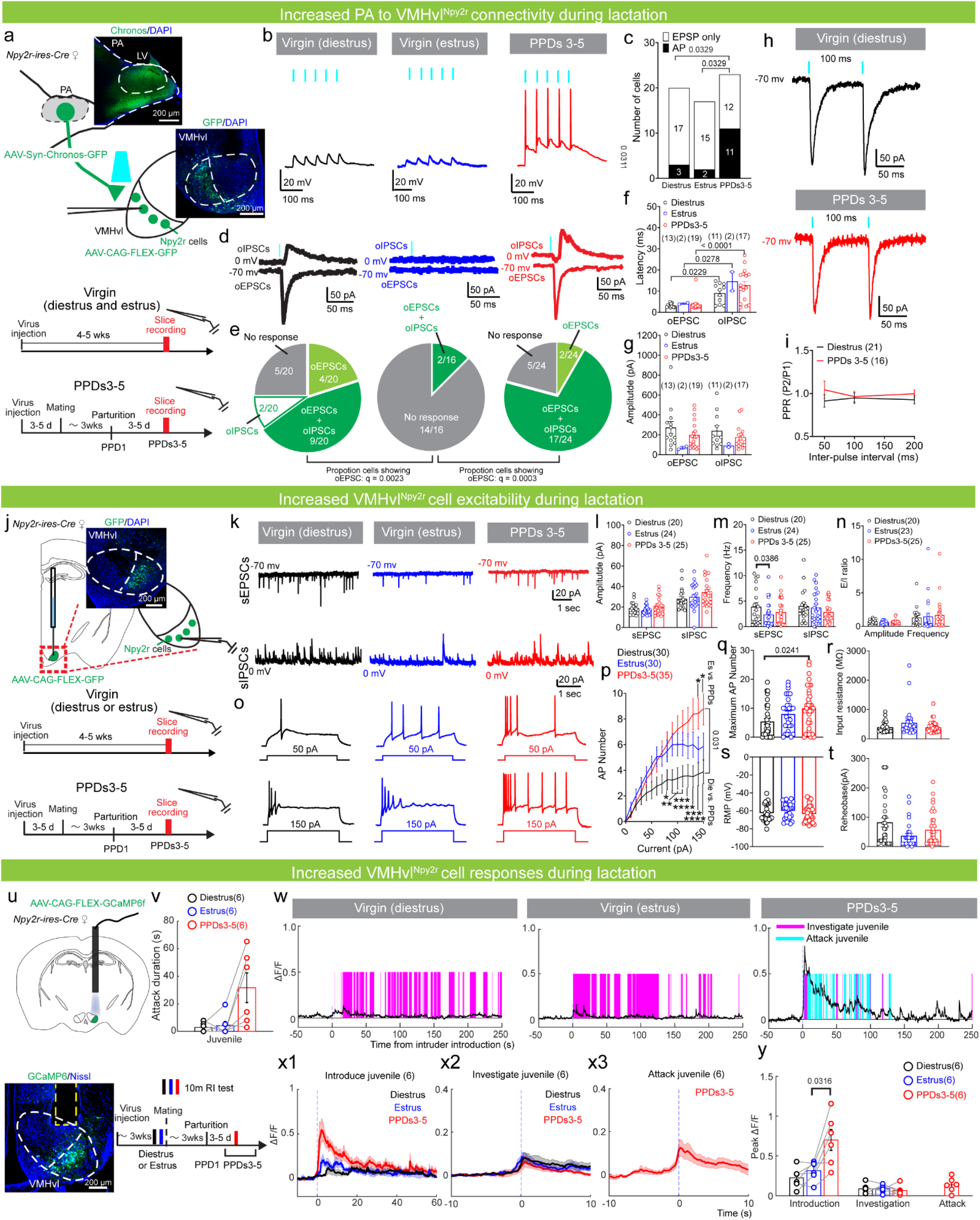
Changes in VMHvl^Npy2r^ responses to intruders and physiology properties in lactating females. **a**, Experimental schematics for the excitatory opsin-assisted circuit mapping and test schedule. Example images showing Chronos-GFP expression in the PA cells (left), and GFP-expressing cells in the VMHvl. Blue, DAPI; scale bar, 200 μm. **b**, Example oEPSP and AP traces evoked by 5-ms light pulses (blue vertical line) in diestrus (black), estrus (blue), and lactating (red) females. **c**, The percentage of cells showing light-induced spiking in each reproductive state. **d**, Example of oEPSC and oIPSC traces evoked by 5-ms light pulses (blue vertical lines) in VMHvl^Npy2r^ cells from diestrous (black), estrous (blue), and lactating (red) females. **e**, Pie charts showing the distribution of light-evoked synaptic responses. **f-g**, The latency (**f**) and amplitude (**g**) of oEPSCs and oIPSCs of VMHvl^Npy2r^ cells in diestrous, estrous, and lactating females. **h**, Examples of oEPSCs of VMHvl^Npy2r^ cells with 100-ms inter-light interval in diestrous and lactating females. **i,** Paired pulse ratios (PPRs) in diestrous and lactating females. **j**, Experimental schematics, test schedule, and a representative image showing GFP (green) expression in the VMHvl^Npy2r^ cells. Blue, DAPI; scale bar, 200 μm. **k**, Example sEPSC and sIPSC traces in diestrous, estrous, and lactating females. **l-n**, The sEPSC and sIPSC amplitude (**l**) and frequency (**m**), and E/I ratio (**n**) of VMHvl^Npy2r^ cells in diestrous, estrous, and lactating females. **o**, Example current-clamp recording traces with 50 and 150 pA current injections of VMHvl^Npy2r^ cells in diestrous, estrous, and lactating females. **p-t**, F-I curves **(p)**, maximum number of APs **(q)**, input resistance **(r)**, RMP **(s),** and rheobase **(t)** of VMHvl^Npy2r^ cells in diestrous, estrous, and lactating females. **u**, Experimental schematics, test schedule, and a representative image showing GCaMP6f (green) expression in VMHvl^Npy2r^ cells. Blue, Nissl; scale bar, 200 μm. **v**, The attack duration of females against juvenile intruders. **w**, Representative Ca^2+^ traces of VMHvl^Npy2r^ cells during interaction with a juvenile intruder in a diestrous (left), estrous (center), or lactating (right) female. The colored shades mark behavioral episodes. **x1-x3**, PETHs of the Ca^2+^ signal of VMHvl^Npy2r^ cells aligned to the intruder introduction (**x1**), investigation (**x2**) and attack onset (**x3**) in the diestrous, estrous or lactating females. **y**, Peak Ca^2+^ signal of VMHvl^Npy2r^ cells during the introduction, investigation, and attack. All bars and error bars and lines and shades represent mean ± s.e.m. Numbers in parentheses indicate the number of recorded cells (**c, e-g, i, l-n, p-t**) or animals (**v, x, y**). Statistical significance was determined using (**c:** Diestrus vs PPDs 3-5, Estrus vs PPDs 3-5, **e**) Chi-square test with FDR correction of Benjamini and Hochberg, (**c:** Diesturs vs Estrus) Fisher’s Exact test with FDR correction of Benjamini and Hochberg, (**f, g, i**) mixed-effects analysis followed by Bonferroni’s multiple-comparison test, (**l-n, q-t**) Kruskal-Wallis test with Dunn’s multiple comparisons, (**p**) two-way ANOVA with repeated measure followed by Bonferroni’s multiple comparison test, (**v**) Friedman test with Dunn’s multiple comparisons, and **(y)** one-way ANOVA with repeated measures followed by Tukey’s multiple-comparisons test. Cells in **(f, g, and i)** were recorded from 5 diestrus, 4 estrus and 6 lactating females. Cells in **(l-m and p-t)** were from 5 diestrous, 4 estrous and 6 lactating females. All p, q values < 0.05 are indicated except (**p**). * < 0.05, ** < 0.01, *** < 0.001, **** < 0.001 in (**p**). See Supplementary Table 1 for more detailed statistics.

Next, we examined the intrinsic properties of VMHvl^Npy2r^ cells using current-clamp recordings. Although rheobase, resting membrane potential (RMP), and input resistance did not differ across reproductive states, VMHvl^Npy2r^ cells in lactating females fired more spikes at nearly all current steps than diestrous females and could reach a significantly higher maximum firing rate (**Fig. 2o-t**). The excitability of VMHvl^Npy2r^ cells in estrous females was between diestrous and lactating females (**Fig. 2o-q**). Collectively, we concluded that the increased spiking probability of VMHvl^Npy2r^ cells to PA input during lactation is likely a combined result of enhanced PA-VMHvl^Npy2r^ connectivity and increased VMHvl^Npy2r^ cell excitability.

The increased VMHvl^Npy2r^ cell responses to inputs can also be observed *in vivo*. As previously reported^10^, Ca^2+^ responses of VMHvl^Npy2r^ cells to juvenile intruders increased significantly during lactation (**Fig 2u-y**). VMHvl^Npy2r^ cells typically show the largest activity increase during initial intruder encounters (**Fig. 2w, x1, y**). In lactating females, the peak Ca^2+^ response during intruder introduction tripled the level in virgin females (**Fig. 2x1, y**). During social investigation, VMHvl^Npy2r^ cells also increased activity consistently, although the response during investigation did not differ between virgin and lactating females (**Fig. 2x2, y**). The females only attacked reliably during lactation (**Fig. 2v**). VMHvl^Npy2r^ cells increased activity consistently and moderately during attack (**Fig. 2x3, y**). Between diestrous and estrous states, VMHvl^Npy2r^ cell Ca^2+^ responses to intruders are similar, consistent with the fact that aggression in virgin females is low regardless of the animal’s estrous state (**Fig. 2v-y**)^9^.

### Limited PA inputs to VMHvl^Npy2r-Esr1+^ cells

PA^Esr1^ cells send dense fibers to both aggression-related VMHvlm and sexual behavior-related VMHvll **(Extended Data Fig. 6a, b).** Yet, we observed no sexual behavior change after inhibiting PA^Esr1→VMHvl^ cells or activating PA^Esr1^ to VMHvl projection (**Extended Data Fig. 3c-i, Extended Data Fig. 4**). To understand why PA input to VMHvl has no impact on female sexual behaviors, we investigated the connection from PA to VMHvl^Npy2r-Esr1+^ cells, a population sufficient to control female sexual receptivity bi-directionally^10,13^. Specifically, we injected AAV-hSyn-Coff/Fon-mCherry into the VMHvl of Npy2r-ires-Cre × Esr1-2A-Flpo mice to label VMHvl^Npy2r-Esr1+^ cells and AAV-hSyn-Chronos-EYFP into the PA to enable optogenetic control of PA to VMHvl terminals on brain slices (**Extended Data Fig. 6c, d**). Strikingly, none of the 18 recorded VMHvl^Npy2r-Esr1+^ cells in estrous females fired a single spike upon PA terminal stimulation, and only 3/15 (20%) cells showed light-evoked EPSCs despite strong Chronos expression in the PA (**Extended Data Fig. 6e-h)**. We focused on estrous females because if PA can influence mating-related VMHvl cell output, it most likely does so during estrus when the mating-related VMHvl cells are most excitable^13^. Indeed, in sexually inactive lactating females, PA exerted even less influence on VMHvl^Npy2r-Esr1+^ cells: only 3/26 (12%) cells showed light-evoked EPSC (**Extended Data Fig. 6g-j**). This directly contrasts the high percentage (79%) of aggression-related VMHvl^Npy2r^ cells showing oEPSC during PA terminal stimulation (**Fig. 2e**). Consistent with the PA terminal stimulation evoked EPSC, the spontaneous sEPSC frequency of VMHvl^Npy2r-Esr1+^ was low during either estrus or lactation compared to VMHvl^Npy2r^ cells (**Extended Data Fig. 6k, l**, **Fig. 2m**).

These results suggest that, first, PA input preferentially targets VMHvl^Npy2r^ cells over VMHvl^Npy2r-Esr1+^ cells, especially during lactation; second, PA inputs could not drive VMHvl^Npy2r-Esr1+^ cells to spike even during estrus, which likely explains the lack of female sexual behavior change during PA^Esr1àVMHvl^ manipulation.

### Increased PA^Esr1^**^→^**^VMHvl^ responses during lactation

Beyond VMHvl cells and PA-VMHvl connection, we wonder whether PA cells also increase responses to aggression-proving cues during lactation to promote aggression. To address this question, we recorded PA^Esr1→VMHvl^ cell calcium activity using fiber photometry (**Fig. 3a**). Specifically, we injected HSV-hEf1α-LS1L-Flpo into the VMHvl and AAV-fDIO-GCaMP6s into the PA of Esr1-2A-Cre virgin female mice with mixed C57 and SW background (**Fig. 3b**). Histology confirms that GCaMP6s is largely confined in the Esr1-expressing cells (92.6%± 0.01% cells, n = 2 animals; **Fig. 3b**). As PA GCaMP6 signal tends to shift over time, likely due to the high expression, we recorded the bulk-calcium activity of PA^Esr1→VMHvl^ cells during virgin and lactating states in separate groups of animals with similar virus incubation periods (**Fig. 3c**). PA^Esr1→VMHvl^ cells increased activity most strongly during intruder introduction, with the highest response during lactation (**Fig. 3e-g)**. During social investigation, PA^Esr1→VMHvl^ cells also responded moderately, and the response magnitude showed a trend of increase in lactating dams (**Fig. 3e, h**). Additionally, PA^Esr1→VMHvl^ cells consistently increased activity during attack, which only occurred in lactating dams (**Fig. 3d-f, i**).

**Figure 3.**
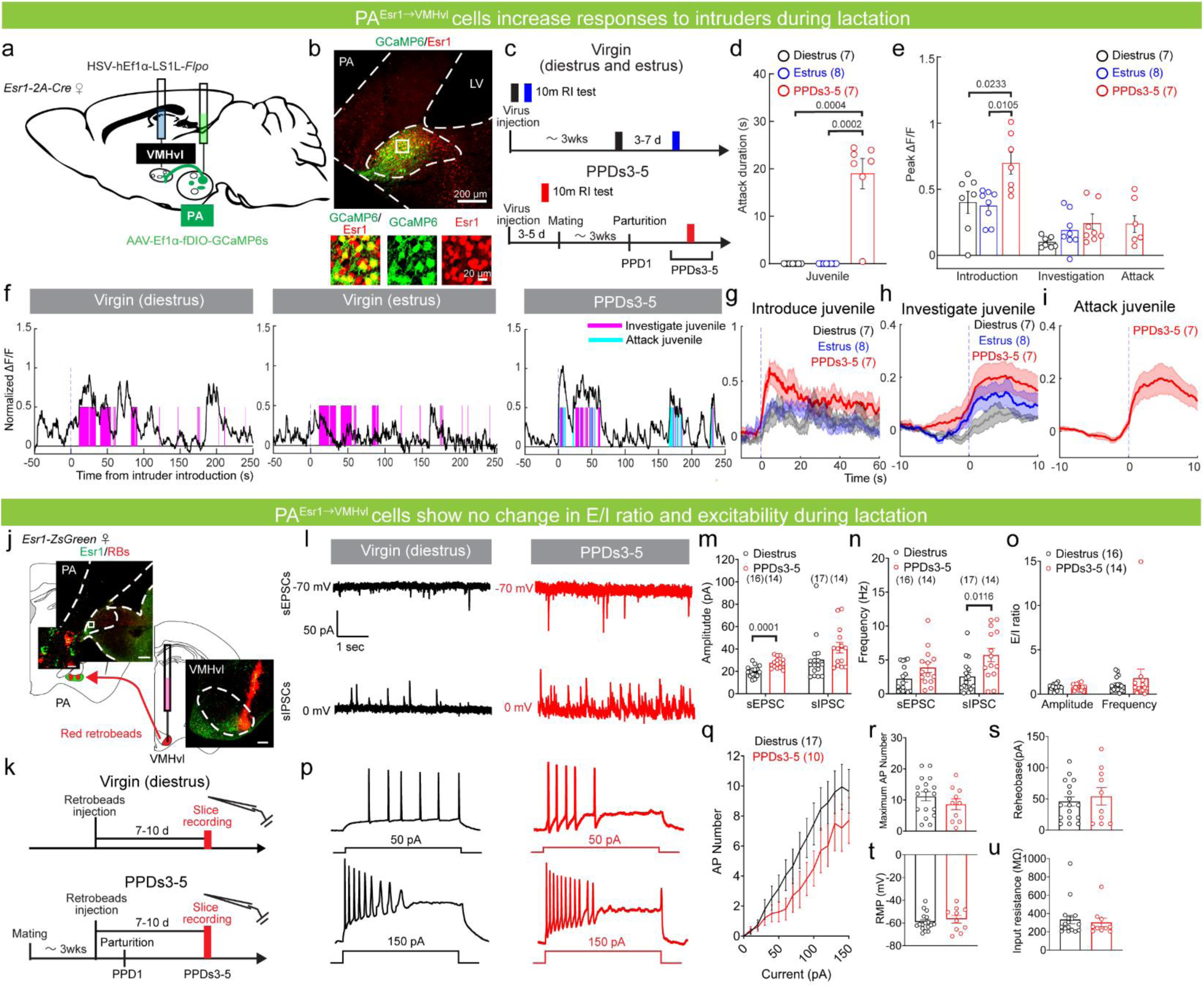
Changes in PA^Esr1 → VMHvl^ responses to intruders and physiology properties in lactating females. **a**, Experimental schematics. **b**, Representative images showing GCaMP6s (green) and Esr1 (red) staining in PA. Bottom: enlarged views of the boxed area; scale bars, 200 μm (top) and 20 μm (bottoms). **c**, Experimental timeline. **d**, The average attack duration of females against juvenile intruders. **e**, Peak Ca^2+^ signal of PA**^Esr1→VMHvl^** cells during the introduction, investigation, and attack. **f**, Representative Ca^2+^ traces of PA**^Esr1→VMHvl^** cells during interaction with a juvenile intruder in a diestrous (left), estrous (center), or lactating (right) female. Color shades mark behavioral episodes. (**g-i**) PETHs of the Ca^2+^ signal of PA^Esr1**→**VMHvl^ cells aligned to the intruder introduction (**g**), investigation (**h**) and attack (**i**) in the diestrous (black), estrous (blue), or lactating (red) females. **j**, Experimental schematics and representative images showing Esr1 expression (green) and VMHvl-injected retrobeads (red) in the PA and VMHvl. Left shows the enlarged view of the boxed area; scale bar, 200 μm (top and bottom) and 20 μm (middle). **k**, Experimental timeline. **l**, Example voltage-clamp recording traces of PA**^Esr1→VMHvl^** cells in diestrous and lactating females. **m-o**, sEPSCs and sIPSCs amplitude **(m)** and frequency **(n)**, and E/I ratio **(o)** of PA **^Esr1→VMHvl^** cells in diestrous and lactating females. **p**, Example current-clamp recording traces with 50 and 150 pA current injections of PA **^Esr1→VMHvl^** cells in diestrous and lactating females. **q-u**, F-I curves **(q)**, maximum AP number **(r)**, rheobase **(s)**, RMP **(t)**, and input resistance **(u)** of PA **^Esr1→VMHvl^** cells in diestrous (black) and lactating (red) females. All bars and error bars and lines and shades represent mean ± s.e.m. Numbers in parentheses indicate the number of animals **(d, e, g-i)** or recorded cells **(m-o, q-u)**. Statistical significance was determined using **(d, e**: Investigation) Kruskal-Wallis test followed by Dunn’s multiple-comparison test, **(e**: Introduction) ordinary one-way ANOVA followed by Tukey’s multiple-comparison test, **(m**: sEPSC, **o**: Amplitude, **r-t)** unpaired t-test, **(m**: sIPSC, **n**: sEPSC and sIPSC, **o**: Frequency, **u)** Mann–Whitney test, and **(q)** two-way ANOVA with repeated measures followed by Bonferroni’s multiple-comparison test. Cells in **(m-o, q-u)** were recorded from 4 diestrus and 4 lactating females. All p values < 0.05 are indicated. See Supplementary Table 1 for more detailed statistics.

To confirm changes in PA^Esr1→VMHvl^ cell responses to the intruder are not due to behavior changes, we recorded the cell responses to various social stimuli in head-fixed awake female mice (**Extended Data Fig. 7a**). Consistent with the response patterns in freely moving animals, PA^Esr1→VMHvl^ cells showed significantly higher responses to juveniles, adult males, and adult females in lactating females than diestrous or estrous virgin females (**Extended Data Fig. 7b-e**). Minimum activity changes of PA^Esr1→VMHvl^ cells were observed during object presentation (**Extended Data Fig. 7b-e**). These in vivo recording data support that PA^Esr1→VMHvl^ cells also undergo changes during lactation, contributing to the overall increase of aggression circuit output that supports the emergence of maternal aggression.

We next performed *in vitro* patch-clamp recordings of PA^Esr1→VMHvl^ cells in brain slices from diestrous virgin and lactating Esr1-Zsgreen C57 female mice to understand which electrophysiological changes may underlie *in vivo* response changes of the cells. We injected retrobeads into the VMHvl and recorded the activity of retrobeads-labeled PA cells in virgin or PPD3-5 females 7-10 days later (**Fig. 3j, k**). Both sEPSC and sIPSC of PA^Esr1→VMHvl^ cells increased moderately during lactation (**Fig. 3l-n**). Specifically, sEPSC amplitude and sIPSC frequency of PA^Esr1→VMHvl^ cells in lactating females were significantly higher than that in virgin females, and sEPSC frequency and sIPSC amplitude also showed trends of increase during lactation (**Fig. 3m, n**). Overall, the excitation-to-inhibition (E/I) ratio did not change during lactation (**Fib. 3o**). Furthermore, unlike the VMHvl^Npy2r^ cells, frequency - current (F-I) curves revealed no difference in PA^Esr1→VMHvl^ cell excitability between lactating and virgin females (**Fig. 3p-r**). Other measurements of intrinsic properties, including rheobase, RMP, and input resistance, also did not differ between virgin and lactating females (**Fig. 3s-u**). These data suggest that PA^Esr1→VMHvl^ and VMHvl^Npy2r^ cells undergo different physiological changes during lactation.

### Oxytocin enhances maternal aggression through PA, not VMHvl

Given a lack of increase in excitability or E/I ratio of PA cells during lactation, we wonder whether additional mechanisms exist to support increased PA^Esr1→VMHvl^ neuron responses to intruders during lactation. Previous studies^22,23^ and gene expression database (mouse.brain-map.org)^24^ suggest highly abundant oxytocin receptor (OXTR) in the PA. Given the high level of oxytocin in lactating mothers and the typical excitatory effect of OXTR via Gq activation^25^, we wonder whether OXTR signaling could also facilitate PA responses during lactation. To address this question, we first examined the OXTR expression in the PA using RNAscope-based *in situ* hybridization assay^26^ and found that virtually all PA^Esr1^ cells express OXTR regardless of the reproductive states (**Fig. 4a-c**). Interestingly, the expression level of OXTR in the PA^Esr1^ cells significantly increased in lactating females, likely indicating increased responses of PA^Esr1^ cells to oxytocin (**Fig. 4a, c**). We next examined the effect of oxytocin on PA^Esr1^ cell activity in lactating females using *in vitro* current-clamp recording (**Fig. 4d**), and found that application of TGOT, a highly specific OXTR agonist, significantly increased PA^Esr1^ cell input resistance but not RMP (**Fig. 4e, f**). In the presence of TGOT, PA^Esr1^ cells fired more action potentials upon 50 pA or 100 pA current injection (**Fig. 4g-i**). Thus, oxytocin can boost the input-output relationship of PA^Esr1^ cells.

**Figure 4.**
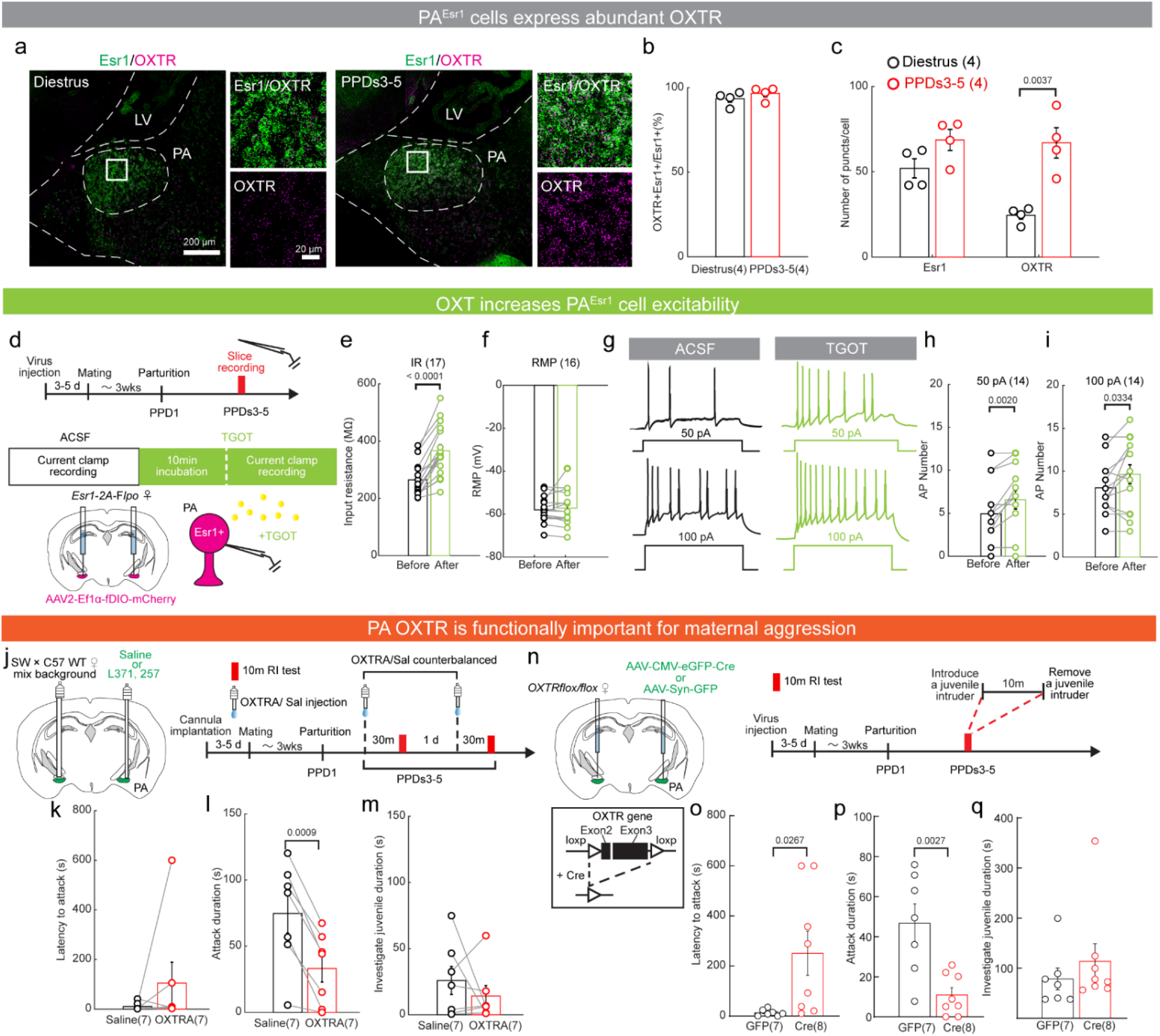
OXTR promotes maternal aggression by enhancing PA activity. **a**, Representative images showing OXTR (magenta) and Esr1 (green) mRNA in the PA in diestrous and lactating females. Right shows the enlarged views of the boxed areas; scale bars, 200 μm (left), 20 μm (right). **b**, The percentage PA^Esr1^ cells expressing OXTR. **c**, Esr1 and OXTR mRNA level per expressing cell in the PA in diestrous and lactating females. **d**, Experimental schematics and timeline. **e, f**, The input resistance **(e)** and RMP **(f)** of PA^Esr1^ cells in lactating females before and after applying TGOT, a specific OXTR agonist. **g**, Example current-clamp recording traces with 50 and 100 pA current injections before and after TGOT application. **h, i**, The spike number of PA^Esr1^ cells with 50 pA **(h)** and 100 pA **(i)** current injections before and after TGOT perfusion. **j**, Experimental schematics and timeline. **k-m**, The latency to attack **(k)**, attack duration **(l)**, and intruder investigation duration **(m)** after injecting saline or L-371,257 hydrochloride, a potent OXTR antagonist (OXTRA). **n**, Experimental schematics and timeline. **o-q**, Latency to attack **(o)**, attack duration **(p),** and intruder investigation duration **(q)** in GFP- and Cre-injected animals. The attack latency was 600 s if no attack was observed. All bars and error bars represent mean ± s.e.m. Numbers in parentheses indicate the number of animals in **(b, c, k-m, o-q)** and recorded cells in **(e, f, h, i)**. Statistical significance was determined using **(b, c, o, p)** unpaired t-test, **(e, k, m)** Wilcoxon matched-pairs signed rank test, **(f, h, i, l)** paired t-test, and **(q)** Mann-Whitney test. Cells in **(e, f, h, i)** were recorded from 4 lactating females. All p values < 0.05 are indicated. See Supplementary Table 1 for more detailed statistics.

To understand the functional importance of PA OXTR signaling in maternal aggression, we injected saline or L-371,257 hydrochloride (100 µM, 250 nl per side), a potent OXTR antagonist (OXTRA), into the PA of lactating female mice 30 min before intruder introduction (**Fig. 4j**). Females attacked the intruder significantly less after OXTRA injection than after saline injection, while the investigation duration and latency to attack did not differ (**Fig. 4k-m**). We also knocked out OXTR in the PA by bilaterally injecting Cre-GFP virus into Oxtr^flox/flox^ female mice (Oxtr^PA-KO^)^27^ (**Fig. 4n**). Control animals were injected with GFP virus (Oxtr^PA-GFP^). Compared to Oxtr^PA-GFP^ females, Oxtr^PA-KO^ females showed significantly shorter attack duration and longer attack latency, while the investigation duration did not change (**Fig. 4o-q**). These functional manipulation results support that OXTR signaling at the PA naturally enhances maternal aggression in lactating females.

OXTR also expresses abundantly in the VMHvl, and similar to PA, the expression level increases during lactation (**Extended Data Fig. 8a, b**). However, we found that the extent of overlap between Npy2r and OXTR is low. Only 28% of Npy2r cells express OXTR in estrous females, which reduces to 18% in lactating females (**Extended Data Fig. 8c, d**). In contrast to PA OXTR manipulation results, injecting OXTRA or conditional knockout of OXTR in the VMHvl caused no significant decrease in attack duration in lactating females, suggesting OXTR signaling does not modulate maternal aggression through VMHvl (**Extended Data Fig. 8e-l**).

### Oxytocin enables pup-dependent adjustment of maternal aggression

As the purpose of maternal aggression is to protect the young, females adjust this costly behavior based on the needs of the young. During the first postpartum week, when the pups are most vulnerable and subjected to infanticide, maternal aggression is the highest (**Fig. 1b-d**)^1,4,5,28^. In scenarios when pups no longer exist, e.g., fall to predation or illness, maternal aggression rapidly decreases^2–4^. Consistent with previous studies^2–4,29^, we found female aggression significantly decreased 24 hours after removing the pups and restored 24 hours after reunion with the pups (**Fig. 5a, c, d**). The serum oxytocin level closely tracked the fall and rise of maternal aggression: it decreased after pup separation and increased after pup reunion (**Fig. 5b**). This is not surprising given that oxytocin is released during nursing, which requires the presence of pups^18,30,31^.

**Figure 5.**
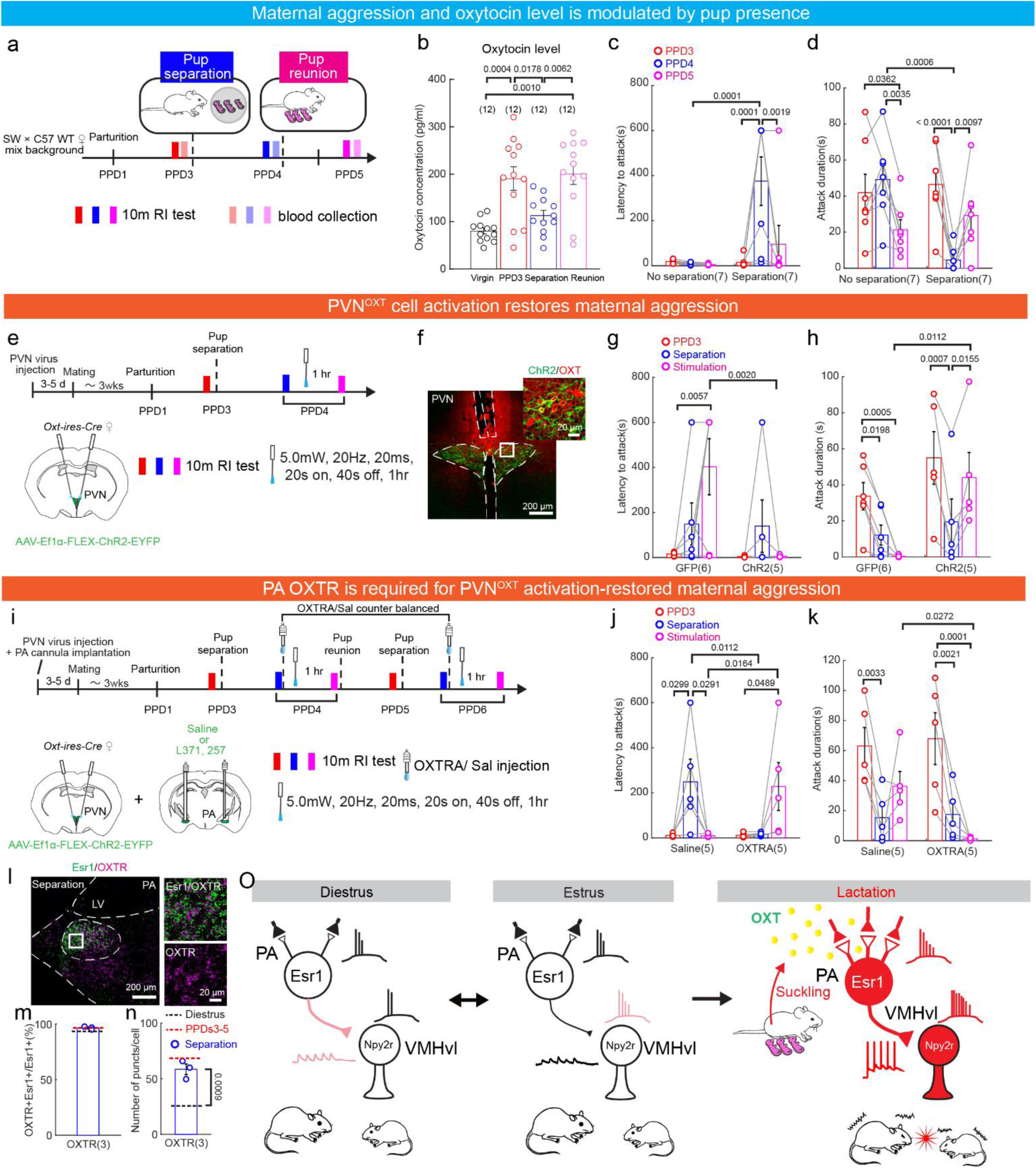
Offspring-dependent oxytocin release is necessary for maintaining maternal aggression. **a**, Experimental schematics to examine the effect of pup separation on maternal aggression. **b**, The serum oxytocin level in virgin females, PPD3 females, PPD4 females 24 hours after pup separation, and PPD5 females 24 hours after pup reunion. **c, d,** The attack latency **(c)** and attack duration **(d)** changes in PPD3 females, PPD4 females 24 hours after pup separation, and PPD5 females 24 hours after pup reunion. The attack latency was 600 s if no attack was observed. **e**, Experimental schematics are used to examine the effect of PVN stimulation in pup separation-caused maternal aggression decrease. **f**, Representative images showing ChR2 (green) expression and OXT (red) staining in PVN. Right shows the enlarged view of the boxed area; scale bars, 200 μm (left) and 20 μm (right). **g, h**, The attack latency **(g)** and attack duration **(h)** in PPD3 females and PPD4 females after pup-separation before and after light delivery to PVN^OXT^ cells in GFP and ChR2 groups. **i**, Experimental schematics to examine the effect of blocking PA OXTR in PVN stimulation-induced maternal aggression recovery after pup separation. **j**, **k**, The attack latency **(j)** and duration **(k)** in PPD3 females and PPD4 females after pup separation before and after light activation to PVN^OXT^ cells in PA OXTRA- and saline-injected groups. **l**, Representative images showing OXTR (magenta) and Esr1 (green) mRNA in the PA in a pup-separated lactating female. Right: enlarged views of the boxed area; scale bars, 200 μm (left), 20 μm (right). **m,n,** The percentage PA^Esr1^ cells expressing OXTR (**m**) and high mRNA levels of OXTR (**n**) in pup-separated lactating females. Black and red dashed lines indicate the average expression levels in virgin and lactating females, respectively. **o,** A model illustrating the neural plasticity underlying the emergence of maternal aggression during lactation. In virgin estrous females, PA-VMHvl^Npy2r^ cell connectivity is weak, while in diestrous females, VMHvl^Npy2r^ cell excitability is low, causing limited spiking of VMHvl^Npy2r^ cells to PA inputs and no aggression towards intruders. In lactating females, PA-VMHvl^Npy2r^ cell connectivity is strong, and VMHvl^Npy2r^ excitability is high, causing increased spiking of VMHvl^Npy2r^ cells to PA inputs. Additionally, the presence of pups, especially the suckling stimuli, induces oxytocin release, further boosting PA^Esr1^ cell output to drive VMHvl^Npy2r^ responses, causing high aggression towards intruders. The synaptic and cellular plasticity at the PA^Esr1^-VMHvl^Npy2r^ pathway enhances the response of the aggression cells to intruders during lactation, while oxytocin enables the lactating female to rapidly adjust its aggression based on the need to protect the young. All bars and error bars represent mean ± s.e.m. Numbers in parentheses indicate the number of animals. Statistical significance was determined using **(b)** Ordinary one-way ANOVA with Tukey’s multiple comparisons test, **(c, d, g, h, j,** and **k)** two-way ANOVA with repeated measures followed by Bonferroni’s multiple-comparison test, and **(m, n)** unpaired t-test. All p values < 0.05 are indicated. See Supplementary Table 1 for more detailed statistics.

Is the decreased maternal aggression after pup separation due to decreased oxytocin? To address this question, we artificially activated oxytocin neurons in the paraventricular hypothalamic nucleus (PVN^OXT^), which are the primary source of oxytocin and are highly activated during pup interaction, especially nursing^30–34^. Specifically, we virally expressed ChR2 (GFP for control) in PVN cells of OXT-Cre virgin female mice (**Fig. 5e, f**). After surgery, the females were paired with males and became mothers. On PPD 3, we tested the female’s aggression level by introducing a juvenile intruder for 10 min. Then, we separated the pups from the females and retested the females’ aggression level 24 hours later (**Fig. 5e**). As expected, both GFP and ChR2 females attacked reliably before pup separation and significantly decreased aggression after pup separation (**Fig. 5g, h**). We then delivered the light (5 mW, 20 Hz, 20 ms, 20 s on and 40s off) to activate PVN^OXT^ cells for 1 hour and retested the aggression level of the female afterward (**Fig. 5e**). Strikingly, the aggression level of ChR2 mice returned to the pre-separation level after light delivery, while GFP mice showed nearly no aggression towards the intruder (**Fig. 5g, h**).

Given the critical role of PA OXTR signaling in boosting female aggression, we asked whether the PVN^OXT^ stimulation-induced recovery of maternal aggression depends on OXTR signaling in the PA (**Fig. 5i**). Before light delivery, we injected either OXTRA or saline into the PA of PVN^OXT^ ChR2 mice and found that OXTRA, but not saline, completely occluded the PVN^OXT^ stimulation-induced maternal aggression recovery (**Fig. 5j, k**). These results suggest that PA OXTR signaling is necessary for PVN^OXT^ to boost maternal aggression rapidly. Importantly, PA OXTR expression one day after pup separation was as high as the pre-separation level, suggesting that PA cells remain highly sensitive to oxytocin (**Fig. 5l-n**). These results suggest that oxytocin, which signals the pup’s presence and development stage, dynamically modulates maternal aggression level to ensure its robust expression but only when necessary.

## Discussion

Female aggression dramatically and abruptly increases after childbirth in mammals to protect offspring. This behavior change must be supported by the increased output of the neural circuit underlying aggression. Our study identified PA^Esr1^ cells as an essential upstream region of the VMHvl^Npy2r^ cells to promote aggression. During lactation, the PA^Esr1^-VMHvl^Npy2r^ circuit shows synaptic and cellular changes, enhancing VMHvl^Npy2r^ cell responses to intruders to drive attack. Furthermore, PA^Esr1^ cells express abundant OXTR, allowing oxytocin to boost the PA cell output. As oxytocin level is tightly linked to the suckling of pups, which varies with the development stage of the pups and, more importantly, the existence of pups, PA^Esr1^ cell output can be rapidly modulated based on the status of the young, ensuring maternal aggression is only expressed to fulfill its protective purpose (**Fig. 5o**).

### Plasticity of PA to VMHvl pathway

The VMHvl has now been firmly established as a key region for social behaviors. In females, two subpopulations in the VMHvl, one expresses Npy2r and the other expresses Cckar, are found important for aggressive and sexual behaviors, respectively^10,12,13^. Our current study on VMHvl^Npy2r^ cells, together with our recent study focusing on VMHvl^Cckar^ cells^13^, revealed their differential connections with PA and distinct changes based on the reproductive state. In virgin females, sexual receptivity is high during estrus and low during diestrus. Consistent with these behavioral characteristics, VMHvl^Cckar^ cells show higher excitability and receive less inhibitory synaptic inputs during estrus than diestrus^13^.

In most laboratory mouse strains, aggression in virgin females is consistently low regardless of their estrous state, yet we noticed some estrous state-dependent physiological changes in the aggression circuit. During diestrus, PA-VMHvl^Npy2r^ connection probability is relatively high, but VMHvl^Npy2r^ cell excitability is low. During estrus, VMHvl^Npy2r^ cell excitability is relatively high, but PA-VMHvl^Npy2r^ connection is nearly absent. The mechanisms underlying these estrous cycle-dependent synaptic and cellular changes remain unclear, but either low excitability or poor synaptic connection makes the aggression circuit difficult to activate in virgin females. This estrous cycle-dependent change in the aggression circuit suggests that female aggression might show cycle-dependent variation in virgin animals. Indeed, wild female mice and female rats are reported to show aggression as virgins, and the aggression level is higher during diestrus than proestrus-estrus^35,36^.

Surprisingly, PA provides nearly no input to the mating-related VMHvl^Npy2r-Esr1+^ cells even during estrus despite the dense fiber terminals in the VMHvll. Behaviorally, PA-VMHvl manipulation could not influence female sexual behaviors in either direction, suggesting that VMHvl mediates female sexual receptivity independent of the excitatory input from PA. Consistent with this hypothesis, sEPSC frequency of VMHvl^Cckar^ and VMHvl^Npy2r-Esr1+^ cells are low, especially when compared to aggression-related VMHvl^Npy2r^ cells (**Fig. 2m, Extended Data Fig. 6l**). The low reliance of mating-related VMHvl cells on excitatory inputs may reflect that female sexual receptivity is largely determined by hormonal state instead of externally driven. Indeed, when female rats are “in heat,” a minimum external stimulus, e.g., touch on the back, is sufficient to induce lordosis^37^.

During lactation, the excitability of Cckar cells is low (maximum firing rate: 5 Hz)^13^, while Npy2r cell excitability becomes high (maximum firing rate: 10 Hz). Furthermore, PA to VMHvl^Npy2r^ cell connection strengthens in that a higher percentage of VMHvl^Npy2r^ cells receive PA input, whereas PA is largely disconnected from VMHvl^Npy2r-Esr1+^ cells. Consequently, PA can effectively drive VMHvl^Npy2r^ cells without engaging mating-related VMHvl cells. This could be critical as mating-related VMHvl cells appear to antagonize aggression-related VMHvl cells^13^. During lactation, when VMHvl^Cckar^ cells are activated, maternal aggression is suppressed even though the female does not show increased sexual receptivity^13^.

### Comparing male and female aggression circuits

PA^Esr1^-VMHvl^Esr1^ circuit is also critical for male aggression. A direct comparison of this circuit between males and females provides important insights into the sex differences in aggression. In males, the majority (∼80%) of VMHvl^Esr1^ cells receive excitatory inputs from PA even when the male is naïve and has no fighting experience. In comparison, 10% and 65% VMHvl^Npy2r^ cells receive PA inputs in estrous and diestrous virgin females. In lactating females, this percentage increases to ∼80%, comparable to naïve males. The connection strength of PA to individual VMHvl cells can readily increase in males, while this appears to be not the case in females. In 5d and 10d male winners, the PA-evoked EPSC amplitude of individual VMHvl^Esr1^ cells is twice that of naïve male mice^15^. In contrast, the PA-evoked EPSC amplitude of individual VMHvl^Npy2r^ cells did not differ between diestrous virgin and lactating females, both similar to the oEPSC amplitude in naïve male mice. In other words, in males, PA and individual VMHvl cells always connect, and the connection strength could increase with winning, whereas in females, the PA to VMHvl connection varies with reproductive states, and the connection strength, if connected, is generally low. Furthermore, the rheobase of VMHvl^Npy2r^ cells in virgin females is higher than that of VMHvl^Esr1^ cells in naïve males^17^, meaning that it takes more excitatory inputs to drive the female aggression-related VMHvl cells to spike. Thus, the larger PA inputs, lower firing threshold, together with the larger number of aggression-related cells in males (recall that only half of female VMHvl^Esr1^ cells are relevant for aggression^9^) make attacks more readily initiated in males than females.

### Oxytocin dynamically signals the need to protect pups

Maternal aggression is unique in that its primary purpose is to protect the young. Thus, the level of maternal aggression needs to track the needs of the young so that females don’t engage in futile fights that are physically demanding and dangerous. Here, we found oxytocin as a critical mediator to link pups’ needs to the aggression circuit output. The high oxytocin level maintained by the pups’ suckling activity is vital to sustaining the high maternal aggression level. Interestingly, PA, not VMHvl, is the key site for oxytocin to boost the aggression circuit output. In the PA, OXTR is broadly and intensely expressed in virtually all PA^Esr1^ cells, enabling oxytocin to increase the input-output relationship of PA^Esr1^ cells by increasing the input resistance of the cell, possibly through closing certain potassium channels^38^. Unlike the role of oxytocin in synaptic plasticity, which could cause lasting changes in the circuit after oxytocin release, the effect of oxytocin on PA cells appears to depend on the presence of the oxytocin. This is likely by design so that oxytocin can couple PA output to pup needs with high temporal precision.

Oxytocin in the PA may originate from direct projections from the PVN^OXT^ cells, as anterograde tracing of PVN^OXT^ cells revealed a few axons in the PA^39^ (mouse.brain-map.org). However, given the sparseness of the axons, we speculate that PA cells may also sense the oxytocin level in the cerebrospinal fluid (CSF), given PA’s close proximity to the lateral ventricle. Indeed, CSF OXT level has been shown to increase during suckling^40–42^. As maternal aggression level does not change from second to second, the slow OXT fluctuation in the CSF should signal pup needs with sufficient temporal resolution to modulate aggression level. The oxytocin-based modulation also has the advantage of being reversible. After pup reunion, oxytocin level quickly rises, and female aggression level returns to the pre-separation level.

As to VMHvl, OXTR signaling may play a role unrelated to female aggression. Our recent study found that oxytocin, originating from the retrochiasmatic supraoptic nucleus (SOR), facilitates social avoidance learning after defeat by enhancing synaptic potentiation of the anterior VMHvl^OXTR^ cells in both male and female mice^43^. Nevertheless, it remains possible that VMHvl oxytocin modulates aggression in males. A recent study showed that mutagenesis of both oxytocin and vasopressin receptors in the VMHvl^Esr1^ cells suppressed male aggression and reduced sustained activity of the VMHvl^Esr1^ cells during inter-male interaction^44^.

In summary, our study revealed neural mechanisms supporting the unique dynamics of female aggression during motherhood. The synaptic and cellular changes, likely induced by the cocktail of sex hormone surges during pregnancy and parturition, primed the aggression circuit to be ready to engage, while oxytocin adds the final layer of control to ensure the aggression is released to serve its purpose of protecting the young. Together, these mechanisms enable the mother to attack the intruder fiercely upon its first encounter while rapidly and reversibly adjusting her aggression level based on the needs of the young.

## Methods

### Mice

All procedures were approved by the NYULMC Institutional Animal Care and Use Committee (IACUC) in compliance with the National Institutes of Health (NIH) Guidelines for the Care and Use of Laboratory Animals. Adult mice aged 12–38 weeks were used for all studies. Mice were housed at 18–23 °C with 40–60% humidity under a 12-h light-dark cycle (dark cycle, 10 p.m. to 10 a.m.), with food and water available ad libitum. For C57 and SW mixed background mice, we bred the F1-mixed background mice by crossing a male C57 Esr1-2A-Cre (Jackson stock no. 017911), Esr1-2A-Flpo (Jackson stock no. 037009), Npy2r-ires-Cre (Jackson stock no. 029285) or Oxt-ires-Cre (Jackson stock no. 024234) with a female SW mouse. For mixed background Npy2r-ires-Cre/Esr1-2A-Flpo double transgenic female mice, we bred the F3-mixed background mice by crossing F2-mixed background homozygous Npy2r-ires-Cre female mice with C57 homozygous Esr1-2A-Flpo male mice. For hybrid OXTR^flox/flox^ female mice (Jackson stock no. 008471), we bred the F2-mixed background mice by crossing F1 SW and C57 mixed background OXTR^flox/+^ mice. These mixed background breeding strategies allowed cell-type-specific manipulations in aggressive lactating female mice. C57 Esr1-ZsGreen mice were generated and kindly provided by Dr. Yong Xu lab originally and then bred in-house. Stimulus animals were adult (>8 weeks) and juvenile (15-25 days old) C57BL/6N female and male mice, originally purchased from Charles River and then bred in-house. Stimulus animals were group-housed. After surgery, all test virgin animals were singly housed, and all lactating animals were co-housed with their pups except during the pup-separation procedure. All experiments were performed during the dark cycle of the animals.

### Viruses

AAV2-CAG-FLEX-GFP (4.00 × 10^12^ vg/mL), AAV2-hSyn-GFP (7.30 × 10^12^ vg/mL), AAV2-Ef1α-DIO-ChR2-EYFP (4.00 × 10^12^ vg/mL), and AAV5-hSyn-Chronos-EYFP (2.10 × 10^13^ vg/mL) viruses were purchased from the University of North Carolina vector core. AAV1-CMV-Hl-eGFP-Cre (2.63 × 10^13^ vg/mL), AAV2-Ef1α-DIO-hM4Di-mCherry (5.00 × 10^12^ vg/mL), AAV2-Ef1α-fDIO-mCherry (8.60 × 10^12^ vg/mL), AAV8-Ef1α-fDIO-GCaMP6s (1.80 × 10^13^ vg/mL) and AAV8-Ef1α-Coff/Fon-mCherry^45^ (2.00 × 10^13^ vg/mL) were purchased from Addgene. AAV2-CAG-FLEX-GCaMP6f-WPRE-SV40 (2.27 × 10^13^ vg/mL) was custom-made by the University of Pennsylvania vector core facility. AAVDJ-Ef1α-fDIO-hM4Di-mCherry (2.65 × 10^13^ vg/mL) was custom-made by Vigene Biosciences, and the plasmid was kindly provided by Dr. Byungkook Lim at USCD. HSV-hEf1α-LS1L-Flpo (1.00 × 10^9^ vg/mL) was purchased from the Massachusetts General Hospital Neuroscience Center vector core.

### Drugs

For chemogenetic inhibition, 5 mg/kg CNO (Sigma, C0832) in saline was intraperitoneally administered. To block OXTRs in the PA and VMHvl, 250 nl per side of 100 µM L-371,257 hydrochloride (Tocris, 2410) in saline was injected through implanted cannulae bilaterally.

### Stereotaxic surgery

Mice (12–20 weeks) were anesthetized with isoflurane (1–1.5%) and placed in a stereotaxic apparatus (Kopf Instruments Model 1900). Viruses and tracers were delivered into the targeted brain regions as described previously^15,46^. Briefly, viruses and tracers were injected into the targeted brain regions using glass capillaries and a nanoinjector (World Precision Instruments, Nanoliter) at 10 and 5 nL/min, respectively. All animals included in the final analysis had correct injection sites, as verified by histology (n = 100 out of 114).

For pharmacogenetic inhibition of PA^Esr1^ cells, 120 nl of AAV2-hSyn-DIO-hM4Di-mCherry was injected bilaterally into the PA (Bregma coordinates: AP, −2.30 mm; ML, ±2.40 mm; DV, −5.80 mm) of Esr1-2A-Cre mice. Test animals that showed correct bilateral targeting of the PA were included in the final analysis (n = 7 out of 8).

For pharmacogenetic inhibition of PA^Esr1→VMHvl^ cells, 130 nl of HSV-hEf1α-LS1L-Flpo was injected into the VMHvl (Bregma coordinates: AP, −1.45 mm; ML, ±0.680 mm, DV, −5.680 mm) while 120 nl of AAVDJ-Ef1α-fDIO-hM4Di-mCherry was injected into the PA bilaterally of Esr1-2A-Cre mice. Control Esr1-2A-Cre animals were bilaterally injected with 130 nl of HSV-hEf1α-LS1L-Flpo into the VMHvl and 100 nl of AAV2-Ef1α-fDIO-mCherry into the PA. Test animals that showed correct bilateral targeting in the VMHvl and virus expression in the PA were included in the final analysis (n = 7 out of 9).

For the optogenetic stimulation of PA-VMHvl projection, 120 nl of AAV2-Ef1α-DIO-ChR2-EYFP was bilaterally injected into the PA of Esr1-2A-Cre mice. Control Esr1-2A-Cre animals were injected with 100 nl of AAV2-CAG-FLEX-GFP bilaterally into the PA. Then, a 200-µm optic-fiber assembly (Thorlabs, FT200EMT, CFLC230) was implanted 200 μm above each side of the VMHvl during the same surgery. Test animals that showed correct bilateral expression of ChR2–EYFP in the PA and correct bilateral fiber placements in the VMHvl were included in the final analysis (n = 7 out of 9).

For the optogenetic stimulation of PVN^Oxy^ cells, 120 nl of AAV2-Ef1α-DIO-ChR2-EYFP was bilaterally injected into the PVN (Bregma coordinates: AP, −0.80 mm; ML, ±0.250 mm, DV, −4.80 mm) of Oxt-ires-Cre mice. Control Oxt-ires-Cre animals were injected with 100 nl of AAV2-CAG-FLEX-GFP into the PVN bilaterally. Then, a 200-µm optic-fiber assembly (Thorlabs, FT200EMT, CFLC230) was implanted 200 μm above each side of the PVN during the same surgery. Test animals that showed correct bilateral expression of ChR2–EYFP or GFP and fiber placement in the PVN were included in the final analysis (ChR2–EYFP: n = 5 out of 6, GFP: n = 6 out of 6).

For combined pharmacological inhibition of PA OXTR and optogenetic stimulation of PVN^Oxy^ cells, 120 nl of AAV2-Ef1α-DIO-ChR2-EYFP was bilaterally injected into the PVN. Then, a 200-µm optic-fiber assembly (Thorlabs, FT200EMT, CFLC230) was placed 200 μm above PVN, and cannulae (RWD Life Science, 62102) were implanted 500 μm above PA bilaterally during the same surgery. All test animals that showed correct bilateral expression of ChR2–EYFP in the PVN and correct fiber and cannulae placements in the PVN and PA were included (n = 6 out of 6).

For fiber photometric recording of the PA^Esr1→VMHvl^ cells, 130 nl of HSV-hEf1α-LS1L-Flpo was injected into the VMHvl, and 120 nl of AAV8-Ef1α-fDIO-GCaMP6s was injected into the ipsilateral PA. Then, a custom-made optic-fiber assembly (Thorlabs, FR400URT, CF440) was inserted ∼150 μm above the dorsal boundary of the PA and secured using dental cement (C&B Metabond, S380). All recordings started 3 weeks after virus injection during virgin and lactating states. Animals that showed correct unilateral targeting in the VMHvl and correct expression of GCaMP6s and fiber placement in the PA were included in the final analysis (n = 16 out of 19).

For fiber photometric recording of VMHvl^Npy2r^ cells, 120 nl of AAV2-CAG-FLEX-GCaMP6f-WPRE-SV40 was injected into the VMHvl of Npy2r-ires-Cre mice. Then, a custom-made optic-fiber assembly (Thorlabs, FR400URT, CF440) was inserted ∼150 μm above the dorsal boundary of the VMHvl and secured using dental cement (C&B Metabond, S380). All recordings started 3 weeks after the virus injection. Animals that showed correct GCaMP6f expression in the VMHvl were included in the final analysis (n = 6 out of 7).

For pharmacological inhibition of OXTR, the cannulae (RWD Life Science, 62102, 62129) were bilaterally implanted 0.5 mm above the PA or VMHvl of mixed background WT mice.

For the conditional gene knockout of OXTR, 200 nl of AAV1-CMV-Hl-eGFP-Cre was injected bilaterally into the PA or VMHvl. Control animals were injected with 100 nl of AAV2-hSyn-GFP bilaterally into the PA or VMHvl. Animals that showed GFP expression in the VMHvl or PA are included in the final analysis (PA-eGFP-Cre: n = 7 out of 9; PA-GFP m = 7 out of 7, VMH-eGFP-Cre: n = 6 out of 6; VMH-GFP: n = 5 out of 5).

For the slice recording of the VMHvl-projecting PA^Esr1^ cells, 30 nl of red retrobeads (Lumafluor Item, R170) was bilaterally injected into the VMHvl of Esr1-zsGreen mice. The brains were used for recording 7-10 days later when the females were either virgin or lactating. Animals that showed correct targeting in the VMHvl were included in the final analysis (n = 4 out of 4).

For the slice recording of the VMHvl^Npy2r^ cells, 100 nl of AAV2-CAG-FLEX-GFP was injected into the VMHvl bilaterally of Npy2r-ires-Cre mice. The brains were used for recording 3-4 weeks later when the females were either virgin or lactating. Animals that showed correct targeting in the VMHvl were included in the final analysis (n = 4 out of 5).

For probing the responses of VMHvl^Npy2r^ and VMHvl^Npy2r-Esr1+^ cells to PA inputs, 200 nl of AAV5-hSyn-Chronos-EYFP was bilaterally injected into the PA, and 200 nl of a 1:1 mixture of AAV2-CAG-FLEX-GFP and AAV8-Ef1α-Coff/Fon-mCherry was bilaterally injected into the VMHvl of Npy2r-ires-Cre × Esr1-2A-Flpo mice. The brains were used for recording 3-4 weeks later when the females were virgin or lactating. Animals that showed correct targeting in VMHvl were included in the final analysis (n = 4 out of 5).

For the slice recording of the PA^Esr1^ cells, 120 nl of AAV2-Ef1α-fDIO-mCherry was bilaterally injected into the PA of Esr1-2A-Flpo mice. The brains were used for recording 3-4 weeks later when the females were virgin or lactating. Animals that showed correct targeting in PA were included in the final analysis (n = 3 out of 3).

### Behavioral analysis

Animal behaviors in all experiments were video recorded from both the side and top of the cage using two synchronized cameras (Basler, acA640-100gm) and a commercial video acquisition software (StreamPix 5, Norpix) in a semi-dark room with infrared illumination at a frame rate of 25 frames per second. Behavioral annotation and tracking were performed frame-by-frame using custom software written in MATLAB (https://pdollar.github.io/toolbox/). For female sexual behaviors, the following definitions applied: ‘lordosis’, the female was on the ground and motionless or showing an arched back posture when the male was mounting or intromitting. The lordosis quotient was calculated as the ratio between lordosis events and male mounting events. The deep thrust success rate was calculated as the ratio between male mount events and the mounting events followed by intromission. For male sexual behaviors, the following definitions were applied: ‘shallowly thrust’, the male posed his forelegs over the female’s back with his hindlimbs on the ground accompanying shallow pelvic thrusts; and ‘deeply thrust’, deep rhythmic thrust following the shallow thrust. For aggression, ‘attack’ was defined as a suite of actions initiated by the resident toward the juvenile intruder, which included lunges, bites, tumbling, and fast locomotion episodes between such behaviors. Investigation was defined by nose contact with any parts of the stimulus animal body. “Pup retrieval” was defined as when the female lifted the pup using her jaw to the moment when the pup was dropped in or around the nest. Mouse behavior was manually annotated frame by frame by an experimenter who was not blind to the group assignment of the animals.

### Female aggression test

To test the female aggression, we performed the resident-intruder test. During the test, we introduced a male or female juvenile intruder mouse (15-25 days old), an adult C57BL/6N male, or an adult C57BL/6N female into the home cage the female for 10 minutes. We did not observe a difference between female aggression towards males and female juvenile intruders. To assess the female aggression level during different reproductive states, we performed the RI tests at approximately the same time the day before mating and on postpartum days 1,3, 5, 7, 14, and 21. When multiple intruders were tested on the same day, the juvenile intruder was always tested first, followed by adult female and adult male intruders. There were at least 2 minutes between the resident-intruder tests. During the postpartum period, the pups were removed 15 minutes before the resident-intruder test and returned after the test.

To examine the effect of pup separation on maternal aggression, all pups were removed from the home cage of the lactating female on PPD3 and cared for by a foster female. 24 hours after pup removal, we introduced a juvenile mouse into the test female’s cage for 10 minutes. After the behavioral testing, all pups were returned to the female’s cage, and 24 hours later, the behaviors towards a juvenile intruder were again examined.

### Fiber photometry

Due to the loss of Ca^2+^ signals in the PA after a long virus incubation period in some animals, we recorded GCaMP6 activity of PA^Esr1→VMHvl^ cells during virgin and lactating states in separate groups of females. In the virgin state, the recording started three to four weeks after the virus injection. Estrous and diestrous states were determined by vaginal cytology. Females were recorded during diestrus and estrus on separate days. During the recording session, we introduced a male or female juvenile (15-25 days old) for 10 minutes. To record the cell responses during lactation, females were co-housed with males three or four days after surgery until the female became visibly pregnant. We then checked the cage daily. The first day when pups were found in the home cage was considered postpartum day 1 (PPD1). The recording in the lactating state was conducted on PPDs 3 to 5, which was typically four to five weeks after virus injection. During the recording session, we removed all the pups and introduced a male or female juvenile (15-25 days old) intruder for 10 minutes.

For the head-fixed recoding test, the mice were habituated to head fixation for at least 3 days, 30 minutes daily. On the day of recording, the mice were head-fixed after RI tests, and an anesthetized male, female, or juvenile C57BL/6N mouse or a 15 ml conical tube was placed approximately 2 mm in front of the nostrils of the recording mouse by an experimenter 4 times, each for 5 s and spaced by 20 s (1 minute between different stimuli).

For fiber photometry recording of the VMHvl^Npy2r^ cells, we monitored Ca^2+^ activity of the same females during virgin and lactating states. Three weeks after the virus injection, the females were recorded during diestrus and estrus. During each recording session, we introduced a male or female juvenile (15-25 days old) for 10 minutes. Afterwards, we paired each female with an adult male and removed the male when the female became visibly pregnant. We then recorded the cell responses to a juvenile intruder for 10 minutes on PPD3-5.

The fiber photometry setup was as described previously^9,15^. Briefly, a 390-Hz sinusoidal blue LED light (30 μW; LED light, M470F1; LED driver, LEDD1B; Thorlabs) was bandpass filtered (passing band, 472 ± 15 nm; FF02-472/30-25, Semrock) and delivered to the brain to excite GCaMP6. The emission lights traveled back through the same optic fiber, were bandpass filtered (passing bands, 535 ± 25 nm; FF01-535/505, Semrock), passed through an adjustable zooming lens (SM1NR01, Thorlabs; Edmund optics no. 62-561), were detected by a Femtowatt Silicon Photoreceiver (Newport, 2151) and were recorded using a real-time processor (RZ5, TDT). The envelope of the 390-Hz signals reflected the intensity of GCaMP6 and was extracted in real-time using a custom TDT OpenEx program. The signal was low-pass filtered with a cutoff frequency of 5 Hz. To analyze the recording data, the MATLAB function ‘msbackadj’ with a moving window of 25% of the total recording duration was first applied to obtain the instantaneous baseline signal. The instantaneous ΔF/F value was calculated as (F_raw_ − F_baseline_)/F_baseline_. The PETH of a given behavior was constructed by aligning the ΔF/F signal to the onset of the behavior. The peak ΔF/F values for each behavior episode were calculated as the maximum ΔF/F 0 to 3s after behavior onset minus the average ΔF/F -10s to -1s before the behavior onset. The introduction peak ΔF/F for each stimulus was calculated as the maximum ΔF/F between 0 and 10 s after the stimulus introduction minus the average ΔF/F -15s to -1s before the behavior onset.

### Pharmacogenetic inactivation

To examine the effect of PA^Esr1^ or PA^Esr1→^ ^VMHvl^ inhibition on female sexual behaviors, we first determined the estrous state of the female using vaginal cytology on the test day. If the female was deemed in estrus, we introduced a highly sexually experienced male into the home cage of the female for approximately 5 minutes to check its behavioral sexual receptivity. The male typically attempted to mount quickly upon encountering the females. If it achieved intromission during the 5-min pretest, the female was considered sexually receptive and proceeded with the test. We first i.p. injected the female with saline. 30 minutes later, we introduced a different sexually experienced male into the female’s home cage for 10 minutes or until the male attempted to mount for 10 times. 1 hour later, we i.p. injected the tested female with 5 mg/kg CNO (Sigma, C0832) we introduced a different sexually experienced male into the female’s home cage again 30 minutes later to re-evaluate the female sexual receptivity. We used different males for the mating tests as the same male would be sexually eager after the first female encounter, which may confound the results. After the test, all females were paired with males until the females became visibly pregnant. We then tested maternal care behaviors and aggression between PPDs 3 and 5. For the maternal care test, all pups except one were removed from the cage immediately after CNO or saline injection. 30 minutes after the injection, we introduced five pups into the female’s home cage far from the nest and observed the female and pup interaction for 10 minutes. Then, we removed all the pups and, two minutes later, introduced a male or female juvenile intruder (15-25 days old) for 10 minutes. One day later, we tested the behaviors of the same animals as the previous day 30 minutes after CNO or saline injection. The CNO or saline injection was counterbalanced across animals.

### Optogenetic activation

After 3 weeks of viral incubation, on the test days, two 200-μm multimode optical fibers (Thorlabs, FT200EMT) were connected with the bilaterally implanted optic fiber assemblies using matching sleeves (Thorlabs, ADAL1). For activating PA^Esr1^ to VMHvl projection in virgin females, on the testing day, we examined vaginal smear 30 minutes before the test to identify diestrous females. During the test, the female mouse was introduced into the home cage of a sexually experienced male. Upon female introduction, the male typically attempted to mount the female quickly. Once the male started to mount frequently and regularly, we delivered 5mW, 20Hz, 20ms, 60s-on, 60s-off light pulses through the bilaterally implanted fibers using a laser (Shanghai Dream Laser) controlled by OpenEx (TDT). The light stimulation was repeated 5-6 times per test session. To test aggressive behaviors, we introduced a male or female juvenile intruder into the home cage of the test female and started the light stimulation (5mW, 20Hz, 20ms, 60s-on, 60s-off) 2 min after juvenile introduction. The light intensity was measured at the fiber end during pulsing using an optical power meter (Thorlabs, PM100D).

For activating VMHvl^Npy2r^ cells in virgin females, 473nm, 20ms, 20Hz light was delivered for 30 s 2-3 minutes after juvenile introduction. The light intensity was 5 mW for real stimulation trials and 0 mW for sham stimulation trials. The order of sham and real stimulation was counterbalanced across animals. Each stimulation type was repeated 4 or 5 times.

To examine the effect of PVN^OXT^ cell activation on maternal aggression, we first tested the baseline aggression 24 hours after pup separation by introducing a male or female juvenile intruder (15-25 days old) into the home cage of the test female for 10 minutes. Then, we delivered 5mW, 20Hz, 20ms, 20s-on, and 40s-off blue light for 1 hour through the implanted optic fibers. Immediately after completing the light stimulation, we again introduced a different juvenile intruder into the female’s home cage for 10 minutes and monitored the behaviors.

### OXTR region-specific knockout

To knock out OXTR in PA and VMHvl, we bilaterally injected 120 nl AAV1 expressing GFP-Cre (control: GFP) into the PA or VMHvl of Oxtr^flox/flox^ female mice. We previously demonstrated that this strategy can successfully knock out OXTR in the VMHvl^43^. Three or four days after surgery, each female was co-housed with a male until the female became visibly pregnant. Between PPD3 and PPD5, we tested the aggression level of the female by introducing a juvenile intruder into the test female’s home cage for 10 min.

### In vivo OXTR antagonist application

To block OXTR in the PA and VMHvl, we bilaterally implanted cannulae (RWD Life Science, 62102, 62129) 0.5 mm above the PA or VMHvl of C57 and SW mixed background female mice. Three or four days after surgery, each female was co-housed with a male until the female became visibly pregnant. Between PPD3 to PPD5, we injected 100 nl 100 µM L-371,257 hydrochloride (Tocris, 2641) into each side of PA or VMHvl through the cannula using a syringe (Hamilton, 65457-02) when the test animal was head-fixed on a running wheel. After the injection, the female returned to its home cage. We introduced a juvenile intruder 30 minutes later and observed the behaviors for 10 minutes. Control animals were injected with saline and underwent the same behavior tests.

### Measure oxytocin

We followed a previous method to measure the oxytocin in the mouse tail blood^47^. Mice were anesthetized with 1.5% isoflurane and placed in a stereotaxic apparatus. We then collected the tail blood and centrifugated it (4 ℃, 1500 RCF) for 10 minutes to isolate the serum. The serum was stored at -80 °C until all samples were collected. We then measured the oxytocin levels in the serum using an ELISA Kit (Arbor Assays, K048-H5) based on the manufacturer’s instructions. We collected blood samples only once from each animal to avoid repeated blood collection-induced changes in maternal behaviors or oxytocin levels.

### In vitro electrophysiological recording

To prepare brain slices for patch-clamp recording, mice were anesthetized with isoflurane, and brains were quickly removed and then immersed in ice-cold cutting solution for 1–2 min (in mM: 110 choline chloride, 25 NaHCO_3_, 2.5 KCl, 7 MgCl_2_, 0.5 CaCl_2_, 1.25 NaH_2_PO_4_, 25 glucose, 11.6 ascorbic acid and 3.1 pyruvic acid). PA or VMHvl coronal sections (275 μm) were cut using a Leica VT1200s vibratome, collected in oxygenated (95% O_2_ and 5% CO_2_) and pre-heated (32– 34 °C) ACSF solution (in mM: 125 NaCl, 2.5 KCl, 1.25 NaH_2_PO_4_, 25 NaHCO_3_, 1 MgCl_2_, 2 CaCl_2_ and 11 glucose) and incubated for 30 min. The sections were then transferred to room temperature and continuously oxygenated until use.

Current and voltage whole-cell patch-clamp recordings were performed using micropipettes filled with intracellular solution containing (in mM) 145 potassium gluconate, 2 MgCl_2_, 2 Na_2_ATP, 10 HEPES, 0.2 EGTA (286 mOsm, pH 7.2) or 135 CsMeSO_3_, 10 HEPES, 1 EGTA, 3.3 QX-314 (chloride salt), 4 Mg-ATP, 0.3 Na-GTP and 8 sodium phosphocreatine (pH 7.3 adjusted with CsOH). Signals were recorded using a MultiClamp 700B amplifier (Molecular Devices) and Clampex 11.0 software (Axon Instruments), and digitized at 20 kHz with Digidata 1550B (Axon Instruments). After recording, data were analyzed using Clampfit (Molecular Devices) or Matlab (Mathworks).

To characterize the physiological and synaptic properties of PA^Esr1^, VMHvl^Npy2r,^ and VMHvl^Npy2r-Esr1+^ cells, we identified ZsGreen, GFP, and mCherry positive cells in the PA or VMHvl on slices from Esr1-ZsGreen mice and Npy2r-ires-Cre mice injected with floxed GFP and Coff/Fon mCherry viruses using an Olympus ×40 water-immersion objective coupled with GFP and TXRED filters. To investigate intrinsic excitability, cells were recorded in the current-clamp mode, and the number of action potentials was counted over 500-ms current steps. The current steps consisted of 18 sweeps from −20 pA to 150 pA at 10 pA per step. sEPSCs and sIPSCs were recorded in the voltage-clamp mode. The membrane voltage was held at −70 mV for sEPSC recordings and 0 mV for sIPSC recordings.

To investigate the synaptic connectivity between PA and VMHvl, Npy2r-ires-Cre × Esr1-2A-Flpo female mice were injected with 200 nl of AAV5-hSyn-Chronos-EYFP into the bilateral PA and 200 nl of a 1:1 mixture of AAV2-CAG-FLEX-GFP and AAV8-Ef1α-Coff/Fon-mCherry into the bilateral VMHvl. Three weeks after virus injection and during recording, we voltage-clamp recorded Npy2r-expressing (GFP positive) or Npy2r-Esr1+ (mCherry positive) VMHvl cells and then delivered 1-ms pulses of full-field illumination using a blue LED light (pE-300white; CoolLED) at 30-s intervals to activate Chronos-expressing PA terminals. oEPSCs and oIPSCs were recorded when holding the membrane potential of recorded neurons at −70 and 0 mV, respectively (corrected for a ∼6 mV liquid junction potential). oEPSPs and APs were recorded under the current clamp mode.

To examine the effect of oxytocin on the cell properties of PA^Esr1^ cells, we injected 120 nl of AAV2-Ef1α-fDIO-mCherry into the PA of Esr1-2A-Flpo female mice. After 3 weeks of virus incubation, we obtained brain slices and performed current-clamp recordings of PA^Esr1^ (mCherry positive) cells. Cell excitability was measured by injecting 500 ms current steps before and after 250 nM TGOT perfusion. The physiological properties of the cells were characterized -10 - 0 min before and 10 - 20 min after TGOT perfusion.

### Immunohistochemistry

Immunofluorescence staining proceeded as previously described^15^. Briefly, mice were perfused transcardially with 1× PBS followed by 4% paraformaldehyde (PFA) in 0.1 M PBS. Brains were extracted, post-fixed in 4% PFA for 3-4 h at 4 °C followed by 48 h in 20% sucrose, embedded in OCT mounting medium, frozen on dry ice, cut to 50-μm-thick sections using a cryostat (Leica Biosystems) and collected in PBS using a 12-well plate. For antibody staining, brain sections were washed with PBS three times and blocked in PBS-T (0.3% Triton X-100 in 1× PBS) with 10% normal donkey serum for 1 h at room temperature. Sections were then incubated in primary antibody diluted in blocking solution at 4 °C for 24–72 h. We stained for Esr1 (rabbit anti-Esr1; 1:1,000; invitrogen, PA1-309, no. YG378288) or Fos (guinea pig anti-Fos; 1:1,000; Synaptic Systems, 226 308). Sections were then washed with PBS-T three times and incubated in secondary antibodies diluted in blocking solution (donkey anti-rabbit, Alexa Fluor 488 or 594, 1:500; Thermo Fisher, A11055 and R37118) with DAPI (1:20,000; Thermo Fisher, D1306) or NeuroTrace 435/455 Blue Fluorescent Nissl Stain (1:200; Thermo Fisher, N21479) for 2–3 h at room temperature. Sections were then washed with PBS three times, mounted on Superfrost slides (Fisher Scientific, 12-550-15), and coverslipped for imaging on a confocal microscope (Zeiss LSM 700 microscope) and/or a virtual slide scanner (Olympus, VS120).

### Fluorescence in situ hybridization

Extracted brains were frozen on dry ice, and 12-μm coronal brain sections were collected using a cryostat (Leica Biosystems). For all genes, we performed RNAscope labeling using Esr1 (478201), Npy2r (315951-C2), and OXTR (412171-C3) probes (Advanced Cell Diagnostics) following the manufacturer’s protocol^26^.

### Cell counting and axon-terminal quantification

To analyze labeled cells, ×20 confocal or epifluorescence images were acquired, and cells were counted manually using ImageJ. Cells that were not entirely contained within a given region of interest were excluded from analyses. For counting DAPI, Nissl, and Esr1-expressing cells, ×20 fluorescence confocal images were acquired. For counting GCaMP6 and mCherry positive cells in pharmacogenetics and fiber photometry experiments, ×20 fluorescence images were acquired with a virtual slide scanner (Olympus, VS120). For counting Esr1, Npy2r, and OXTR positive cells, ×20 fluorescence confocal images were acquired. To quantify Esr1 and OXTR expression at the single-cell level, the number of puncta in individual cells was counted using ImageJ. To quantify the density of of PA^Esr1^ cell axons in the VMHvl, we selected a boxed area (200 × 200 μm^2^) in each subregion containing fibers and calculated the average pixel intensity as *F*_raw_ using ImageJ and Photoshop. On the same image, a boxed area of the same size in the dorsomedial part of VMH (VMHdm) containing no fiber terminals was selected to calculate the background intensity (*F*_background_). *F*_signal_ was then calculated as *F*_raw_ – *F*_background_. For each animal, *F*_signal_ was normalized by the maximum *F*_signal_ across all the analyzed regions. The normalized *F*_signal_ was then used to calculate the average terminal field intensity across animals.

### Statistics

All statistical analyses were performed using MATLAB2020a (Mathworks) or Prism7 software (GraphPad). All datasets were tested for normality with the Shapiro-Wilk test, except for repeated-measures two-way ANOVA, mixed-effects analysis, Cochran’s Q test, McNemar test, Chi-square test, and Fisher’s exact test. If the dataset passed the normality test, parametric tests were used, including unpaired t-test, paired t-test, repeated-measures, and ordinary one-way ANOVA with Geisser–Greenhouse correction and Tukey’s multiple-comparison post hoc test. Otherwise, non-parametric tests were used, including Mann–Whitney test, Wilcoxon matched-pairs signed-rank test, Friedman test with Dunn’s multiple-comparison test, Kruskal-Wallis test with Dunn’s multiple comparisons test. Repeated measures and ordinary two-way ANOVA with Bonferroni’s multiple-comparison post hoc test were used for statistical tests containing two variables. Mixed model analysis was used to analyze repeated measures data when there were missing values. Following repeated-measures two-way ANOVA and mixed-effect model, post hoc Bonferroni’s multiple-comparison test was performed. Cochran’s Q test, Chi-square test, McNemar test, and Fisher’s exact test were used for categorical data. When multiple pairwise tests were performed, the p values were adjusted using the false discovery rate (FDR = 0.05). All statistical tests are two-sided. All significant statistical results (p < 0.05) were indicated in the figures. All error bars indicate s.e.m. No statistical methods were used to predetermine sample sizes, but our sample sizes were similar to those reported previously^15,46^. See Supplementary Table 1 for more statistical details.

## Acknowledgments

We thank all of the members from the Lin laboratory for the discussions, Yiwen Jiang for assistance with genotyping, Dr. Xu Yong for sharing Esr1-zsGreen mouse, Dr. Stephen Liberles for sharing Npy2r-ires-Cre mouse, Dr. Karl Deisseroth & INTRSECT 2.0 Project for providing pAAV-Ef1a-Con/Fon-mCherry plasmid and Dr. Byungkook Lim for providing AAV-Ef1α-fDIO-hM4Di-mCherry plasmid. We thank Long Mei for assistance with brain atlas images.

## Author contribution

D.L. and T.Y. conceived the project, designed the experiments, and wrote the paper. D.L. supervised the projects. T.Y. performed all functional manipulation, *in vivo* recording, and histology experiments and analyzed data. R.Y. performed all *in vitro* slice recording experiments, analyzed data, and wrote the relevant results. M.K. and K.T. assisted with behavioral testing and annotation. T.O. performed preliminary testing of the VMHvl OXTR role in maternal aggression. S.P. and N.M.S. generated Esr1-2A-Flpo knock-in mice.

---

**Extended Data Figure 1.**
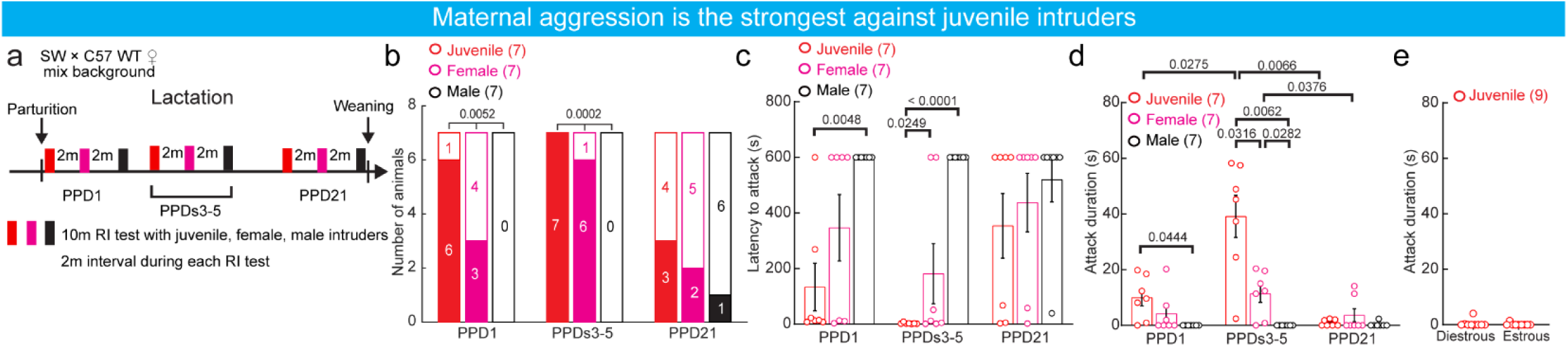
The level of maternal aggression towards different intruders. **a,** Experimental timeline. **b**, The number of C57 × SW females that attacked a male, female, or juvenile intruder during different lactation periods. **c,** The latency to attack a juvenile, an adult female, and an adult male intruder in mothers at different lactation periods. **d,** The duration of attack against a juvenile, an adult female, and an adult male intruder in mothers at different lactation periods. **e**, No virgin females attacked a juvenile intruder regardless of their estrous states. All bars and error bars represent mean ± s.e.m. Numbers in parentheses indicate the number of animals. Statistical significance was determined using **(b)** Chi-square test followed by Mcnemar test with FDR correction of Benjamini and Hochberg, **(c, d)** two-way ANOVA with repeated measures followed by Bonferroni’s multiple-comparison test, and **(e)** Mann-Whitney test. All p values < 0.05 are indicated. See Supplementary Table 1 for more detailed statistics.

**Extended Data Figure 2.**
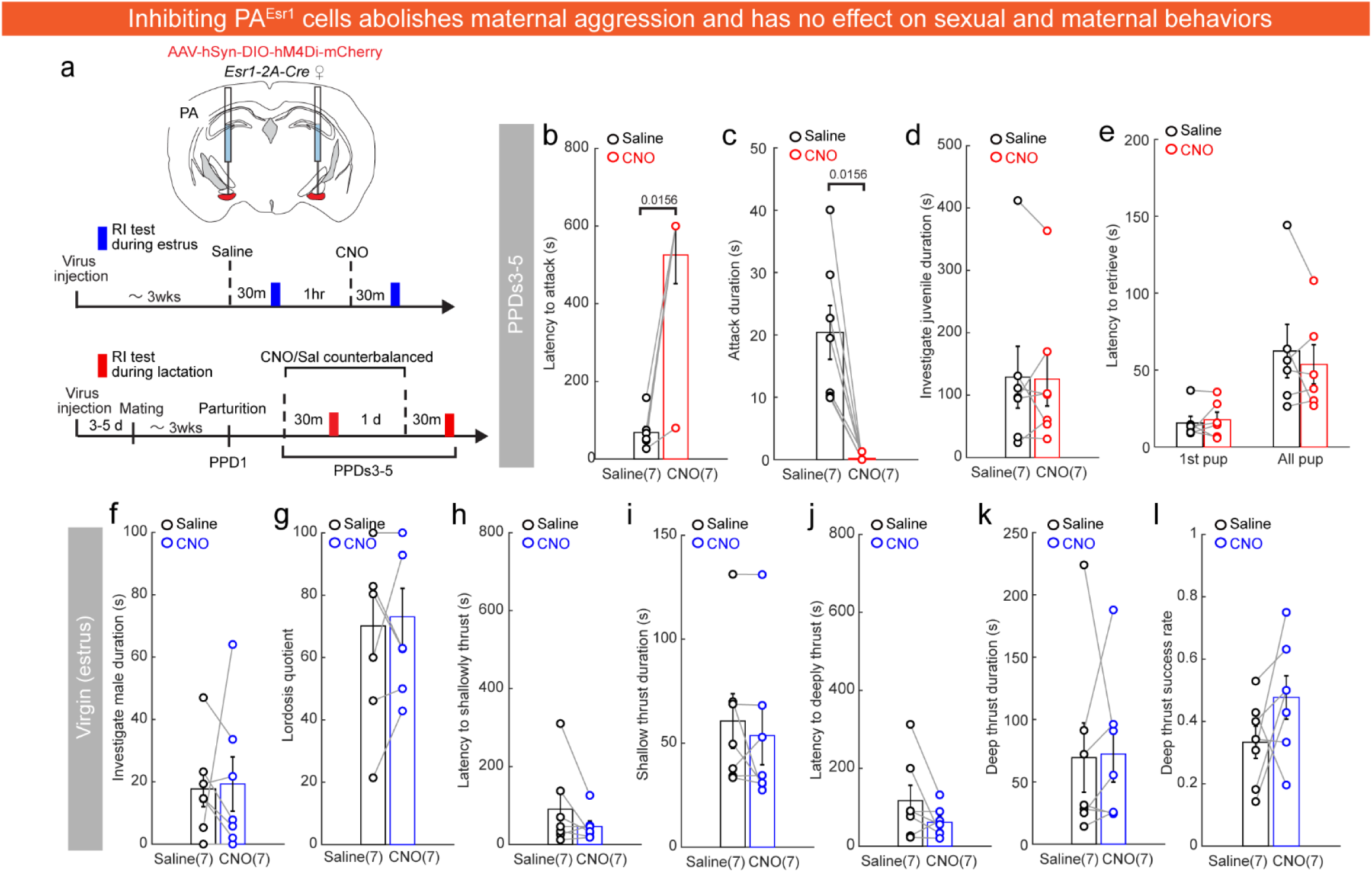
Chemogenetic inhibition of PA^Esr1^ cells impairs maternal aggression, not maternal care or sexual receptivity. **a,** Experimental schematics and timeline. **b-d**, The latency to attack (**b**), attack duration (**c**), and time spent investigating juvenile intruders **(d)** after saline or CNO injections in lactating females that expressed hM4Di in PA^Esr1^ cells. **e**, The latency to retrieve the first pup and all 5 pups after saline or CNO injections in lactating females that expressed hM4Di in PA^Esr1^ cells. **f, g**, The male investigation duration **(f)** and lordosis quotient **(g)** after saline or CNO injections in virgin estrous females that expressed hM4Di in PA^Esr1^ cells. **h-l**, The latency to achieve shallow thrust **(i),** duration of shallow thrust **(j),** latency to deep thrust **(k),** deep thrust duration **(l),** and deep thrust success rate **(l)** when sexually experienced males mated with estrous females that expressed hM4Di in PA^Esr1^ cells and received either CNO or saline injection. All bars and error bars represent mean ± s.e.m. Numbers in parentheses indicate the number of animals. Statistical significance was determined using **(b-d, e**:1^st^ pup**, h-j)** Wilcoxon matched-pairs signed-rank test and **(e: all pup, f, g, k, l)** paired-t-test. All p values < 0.05 are indicated. See Supplementary Table 1 for more detailed statistics.

**Extended Data Figure 3.**
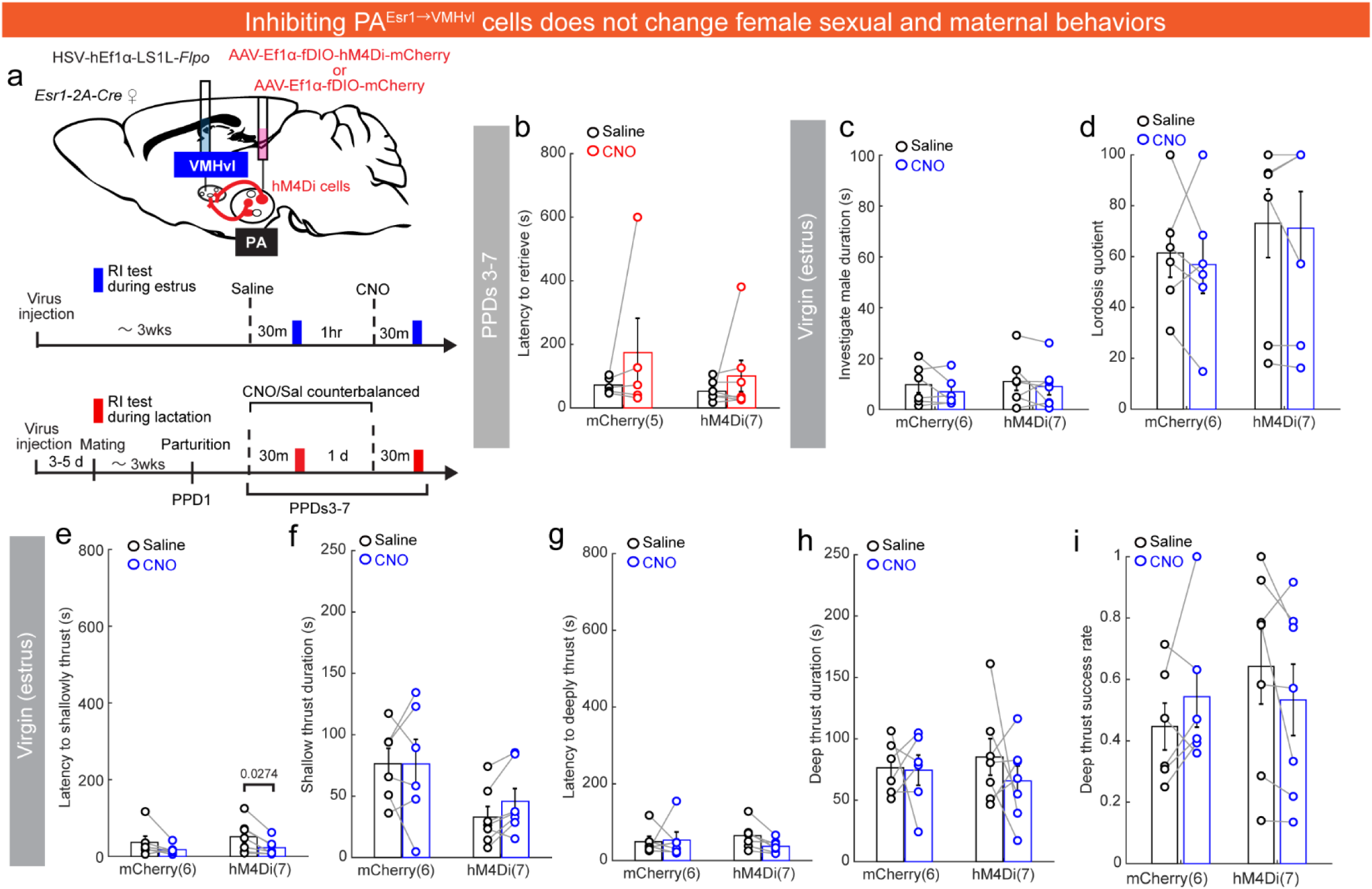
Chemogenetic inhibition of PA^Esr1 → VMHvl^ cells does not change sexual receptivity or maternal care. **a,** Experimental schematics or timeline. **b**, The latency to retrieve the first pup and all 5 pups after saline or CNO injections in lactating females that expressed hM4Di in PA^Esr1→VMHvl^ cells. The latency was 600 s if no retrieve occurred. **c, d,** The male investigation duration **(c)** and lordosis quotient **(d)** after saline or CNO injections in virgin estrous females that expressed hM4Di in PA^Esr1→VMHvl^ cells. **e-i**, The latency to achieve shallow thrust **(e),** duration of shallow thrust **(f),** latency to deep thrust **(g),** deep thrust duration **(h)**, and deep thrust success rate **(i)** when sexually experienced males mated with estrous females that expressed hM4Di in PA^Esr1→VMHvl^ cells and received either CNO or saline injection. All bars and error bars represent mean ± s.e.m. Numbers in parentheses indicate the number of animals. Statistical significance in **(b-i)** was determined using two-way ANOVA with repeated measures followed by Bonferroni’s multiple-comparison test. All p values < 0.05 are indicated. See Supplementary Table 1 for more detailed statistics.

**Extended Data Figure 4.**
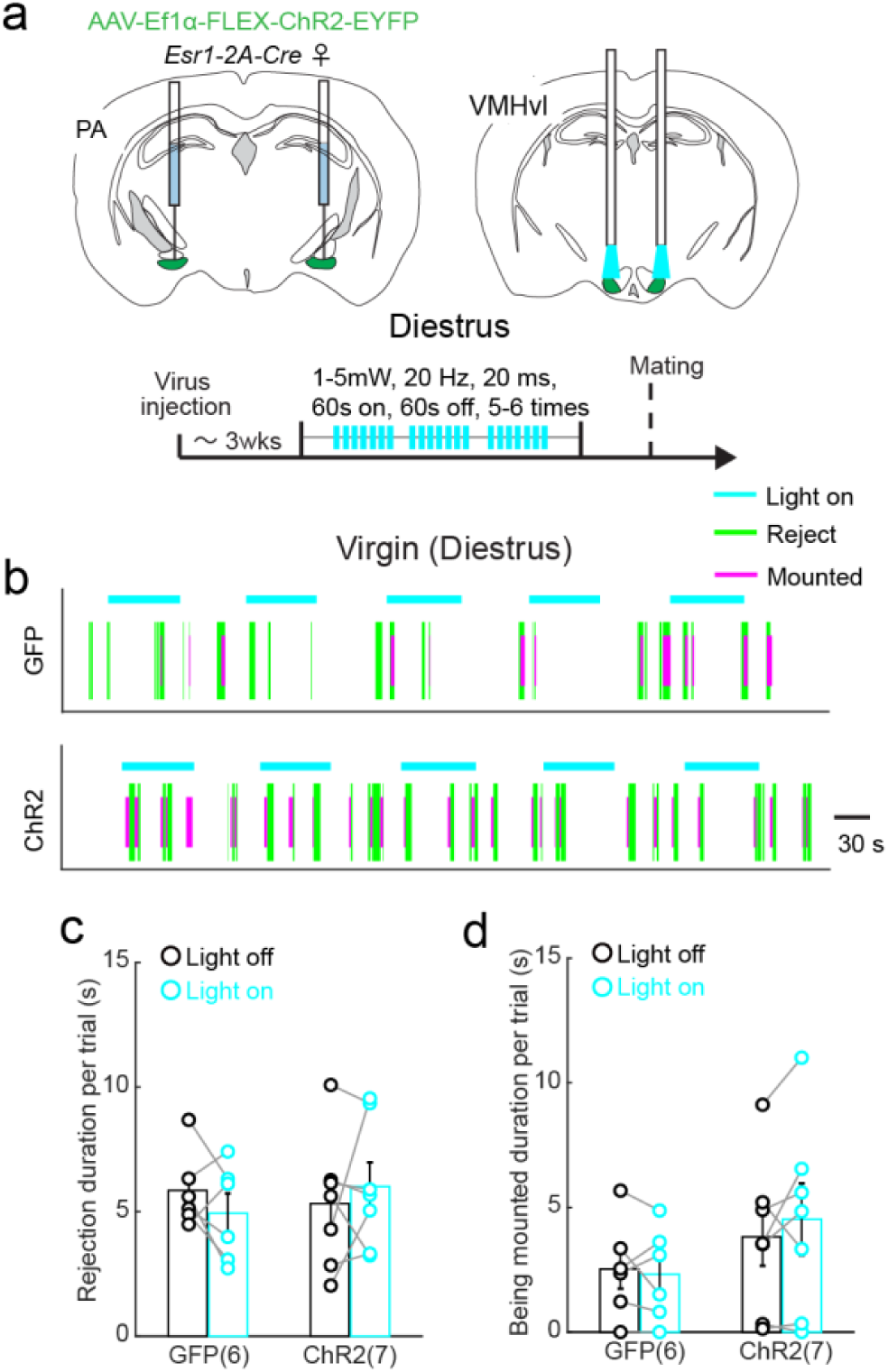
Optogenetic activation of PA^Esr1^ to VMHvl projection does not change female receptivity. **a,** Experimental schematics and timeline. **b**, Representative raster plots showing behaviors of virgin diestrous females expressing ChR2 in PA^Esr1^ cells during interaction with a male intruder when blue light was delivered to the VMHvl. Top showing light-on period in cyan. Bottom showing behaviors. **c, d**, The time females spent rejecting the male **(c)**, and the duration of male mounting **(d)** during light on and light off periods. All bars and error bars represent mean ± s.e.m. Numbers in parentheses indicate the number of animals. Statistical significance in **(c, d)** was determined using two-way ANOVA with repeated measures followed by Bonferroni’s multiple-comparison test. All p values < 0.05 are indicated. See Supplementary Table 1 for more detailed statistics.

**Extended Data Figure 5.**
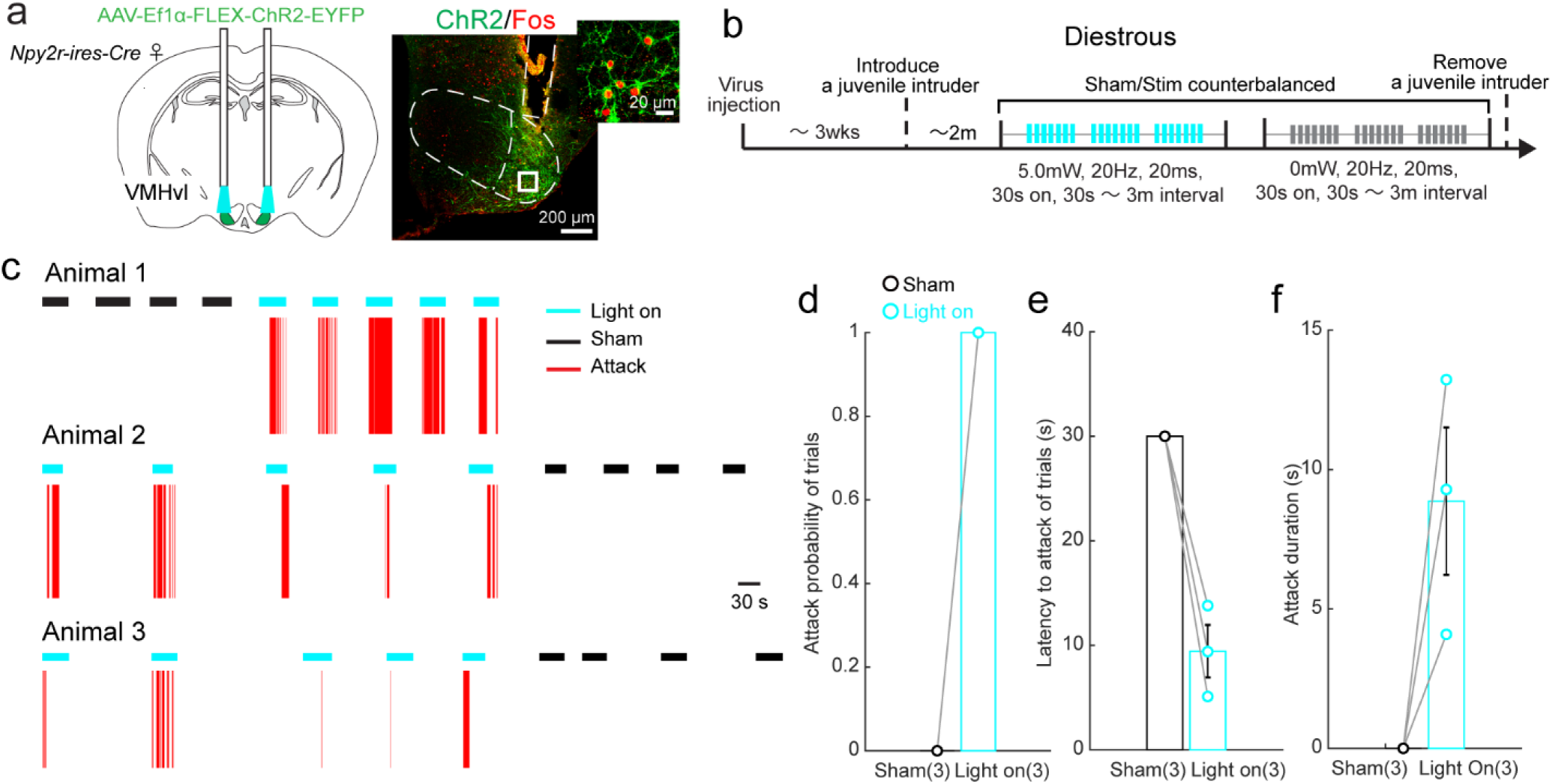
Optogenetic activation of VMHvl^Npy2r^ cells induced time-locked attack in virgin females. **a**, Experimental schematics and representative images showing ChR2 (green) and Fos (red) expression in the VMHvl. Right shows the enlarged view of the boxed area; scale bars, 200 μm (left) and 20 μm (right). **b**, Experimental timeline. **c**, Representative raster plots showing behaviors of virgin females that expressed ChR2 in VMHvl^Npy2r^ cells during interaction with a juvenile intruder when blue light or sham light was delivered to the VMHvl. Top showing sham (black) or light (blue) delivery periods. Bottom showing behaviors. **d-f**, The proportion of trials that animals attacked **(d)**, the latency to attack during each trial **(e)**, and the average attack duration per trial **(f)** towards juvenile intruders of light-on and sham trials. Bars and error bars represent mean ± s.e.m. Numbers in parentheses indicate the number of animals. Statistical significance in **(d-f)** was determined by Wilcoxon matched-pairs signed-rank test. All p values < 0.05 are indicated. See Supplementary Table 1 for more detailed statistics.

**Extended Data Figure 6.**
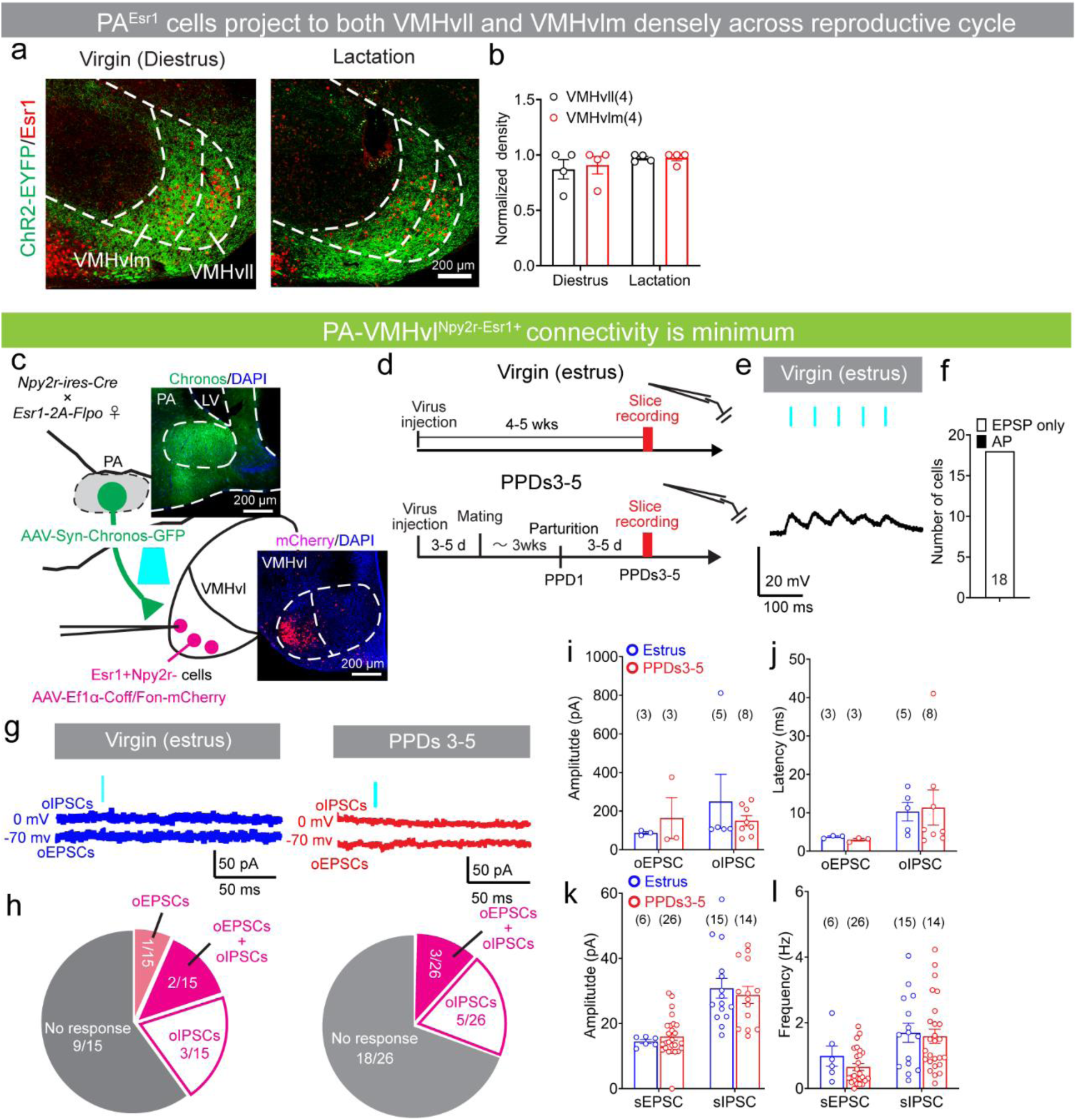
PA to VMHvl^Npy2r-Esr1+^ cell connectivity in virgin and lactating females. **a**, Representative images showing terminals from PA^Esr1^ cells expressing ChR2-EYFP in the medial and lateral subdivisions of the VMHvl (VMHvlm and VMHvll) of diestrous and lactating females. Scale bar, 200 μm. **b**, The normalized density of PA^Esr1^ cell terminals in VMHvlm and VMHvll in diestrous and lactating females. *F*_raw_ was calculated the average pixel intensity in each subregion containing fibers and dorsal medial part of VMH containing no fiber terminals was selected for calculating the background intensity (*F*_background_). *F*_signal_ was then calculated as *F*_raw_ – *F*_background_. For each animal, *F*_signal_ was normalized by the maximum *F*_signal_ across all the analyzed regions. The normalized *F*_signal_ was then used for calculating the average terminal field density across animals. **c**, Experimental schematics for the excitatory opsin-assisted circuit mapping. Example images showing Chronos-GFP expression in the PA and mCherry expression in the VMHvl. Scale bars, 200 μm. **d**, Experimental timeline. **e**, Example voltage clamp recording trace of a VMHvl^Npy2r-Esr1+^ cell in response to 5-ms blue-light pulses (blue vertical lines). **f,** The percentage of VMHvl^Npy2r-Esr1+^ cells that showed PA terminal stimulation-evoked spiking in estrous females. (n = 18 cells from 3 estrus females). **g**, Example voltage-camp recording traces upon 5-ms blue-light pulses (blue vertical lines) of VMHvl^Npy2r-Esr1+^ cells in estrous (left) and lactating (right) females. **h,** Pie charts showing light-evoked synaptic response patterns of VMHvl^Npy2r-Esr1+^ cells in estrous (left) and lactating (right) females. **i, j**, The amplitude **(i)** and latency **(j)** of oEPSCs and oIPSCs of VMHvl^Npy2r-Esr1+^ cells in estrous and lactating females. The cells were recorded from 3 estrous and 6 lactating females. **k, l,** The amplitude **(k)** and frequency **(l)** of sEPSCs and sIPSCs of VMHvl^Npy2r-Esr1+^ cells in estrous and lactating females. All bars and error bars represent mean ± s.e.m. Numbers in parentheses indicate the number of recorded cells. Statistical significance was determined using **(b)** two-way ANOVA with repeated measures followed by Bonferroni’s multiple-comparison test, **(h)** chi-square test, **(i, j)** Mixed-effects analysis with Bonferroni’s multiple comparisons test, and **(k, l)** Mann-Whitney test. The cells were recorded from 3 estrous and 6 lactating females. All p values < 0.05 are indicated. See Supplementary Table 1 for more detailed statistics.

**Extended Data Figure 7.**
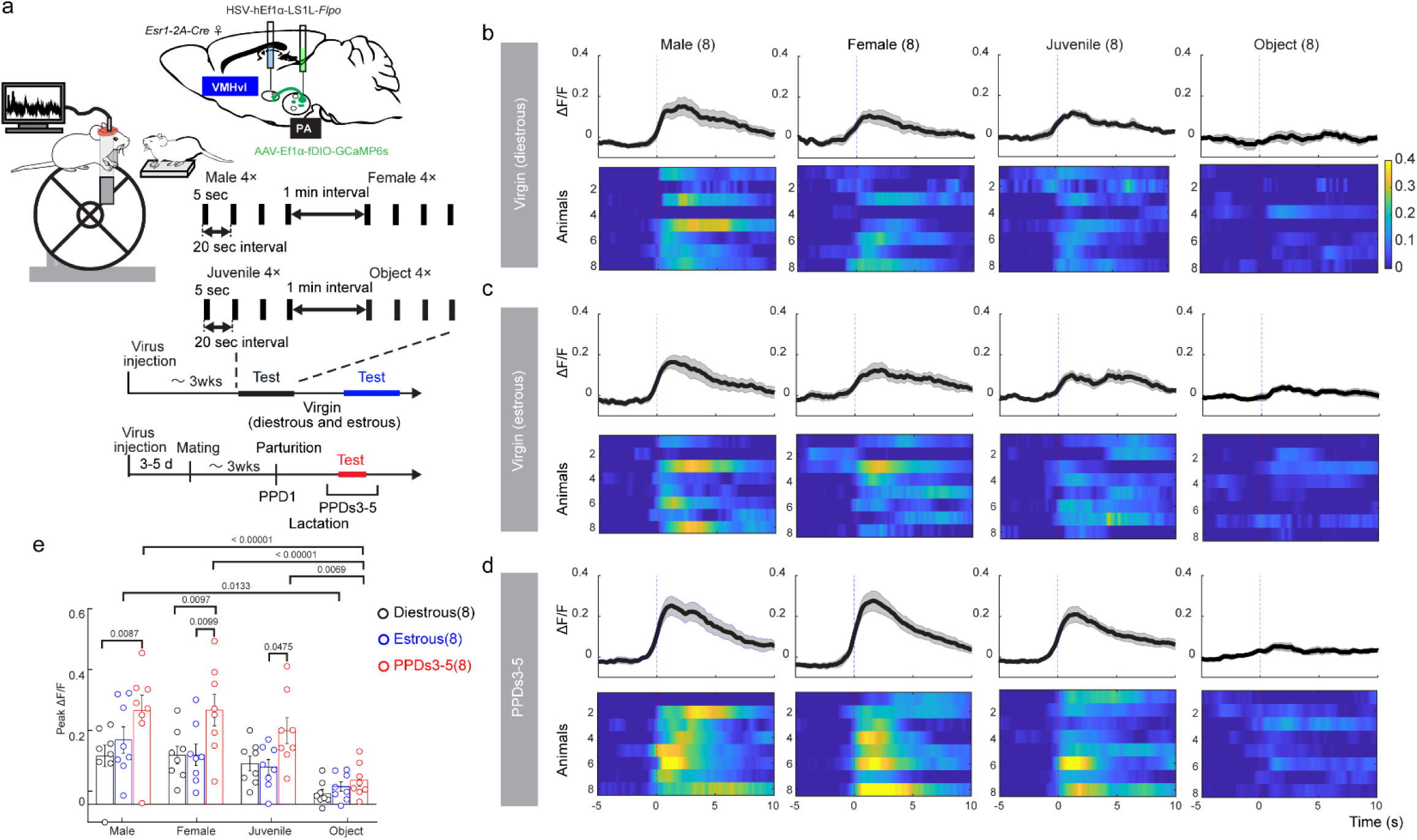
Female PA^Esr1→VMHvl^ cells show higher responses to social cues during lactation. **a**, Experimental schematics for head-fixed recording and experimental timeline. **b-d**, PETHs (top) and heatmaps (bottom) of PA^Esr1→VMHvl^ cell Ca^2+^ signals aligned to the onset of adult male, adult female, juvenile, and object presentations when the recording females were in diestrus (**b**), estrus (**c**), and lactation (**d**). **e**, Peak ΔF/F during the presentation of various stimuli when the recording females were in diestrus, estrus, and lactation. All data are presented as the mean ± s.e.m. Numbers in parentheses indicate the number of recording mice. Statistical significance in **(e)** was determined using ordinary two-way ANOVA followed by Bonferroni’s multiple-comparison test. All p values < 0.05 are indicated. See Supplementary Table 1 for more detailed statistics.

**Extended Data Figure 8.**
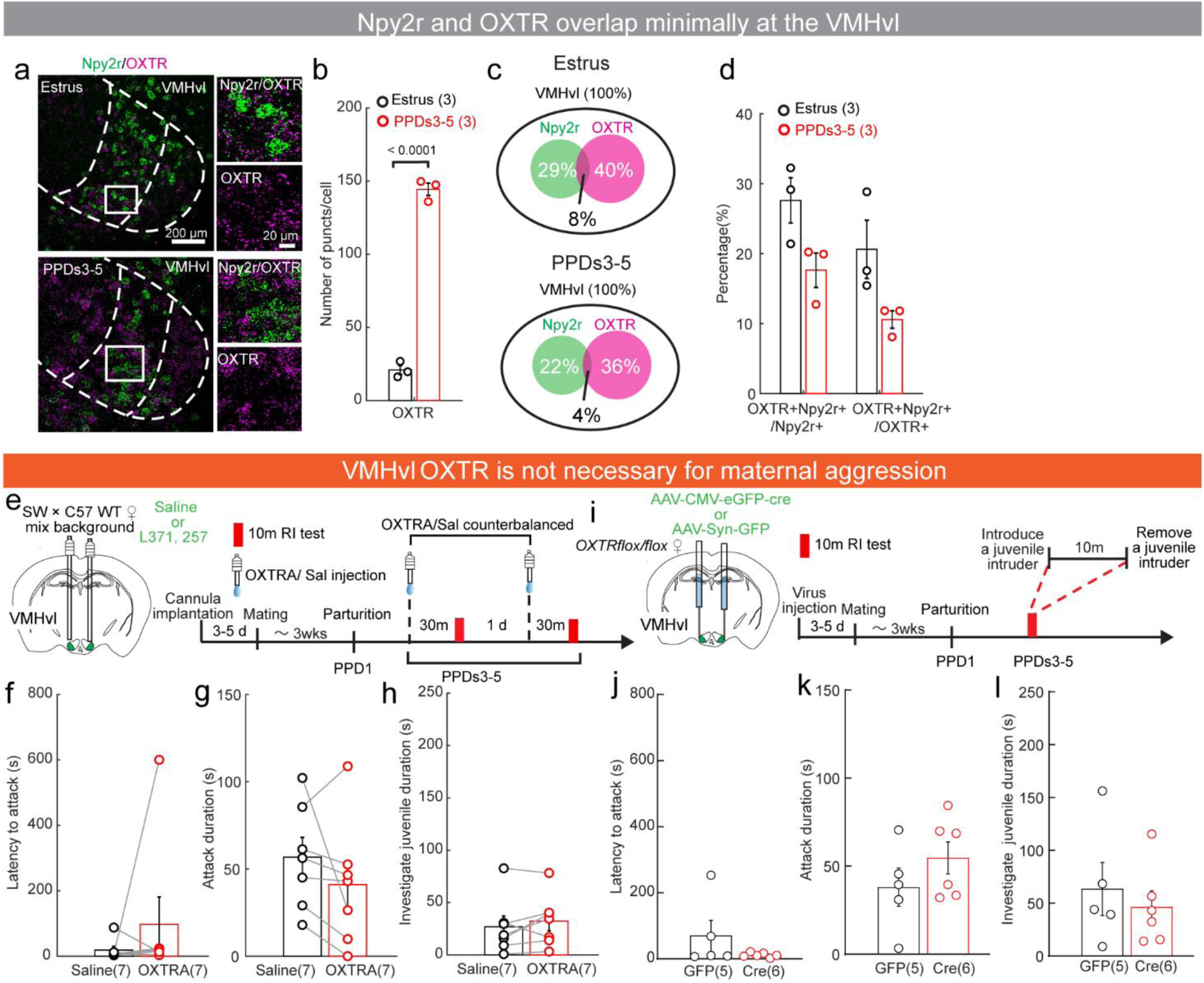
VMHvl OXTR signaling is not essential for maternal aggression. **a**, Representative images showing OXTR (magenta) and Npy2r (green) mRNA in the VMHvl in estrous and lactating females. Right shows the enlarged views of the boxed area; scale bars, 200 μm (left), 20 μm (right). **b**, The number of OXTR mRNA puncts per OXTR-expressing cell in the VMHvl of estrous and lactating females. **c**, Venn diagrams showing the percentage of Npy2r and OXTR positive cells in the VMHvl in estrous (top) and lactating (bottom) females. **d**, The percentages of Npy2r and OXTR double-positive cells in NPY2r or OXTR expressing cells. **e**, Experimental schematics and timeline to examine the effect of OXTR antagonist in the VMHvl in maternal aggression. **f-h**, The latency to attack **(f),** attack duration **(g),** and juvenile investigation duration **(h)** did not differ after OXTRA (red) and saline (black) injections. The attack latency was 600 s if no attack occurred. **i**, Experimental schematics and timeline to examine the effect of VMHvl OXTR knockdown in maternal aggression. **j-l**, Conditional OXTR knockout in the VMHvl did not change latency to attack **(j),** attack duration **(k)**, and juvenile investigation duration **(l)**. All data are presented as the mean ± s.e.m. Numbers in parentheses indicate the number of subject mice. Statistical significance was determined using **(b, d**:Npy2r+OXTR/Npy2r**, k)** unpaired t-test, **(d**: Npy2r+OXTR/OXTR**, j, l)** Mann-Whitney test, **(f)** Wilcoxon matched-pairs signed rank test, and **(g, h)** paired t-test. All p values < 0.05 are indicated. See Supplementary Table 1 for more detailed statistics.

